# Light-inducible Deformation of Mitochondria in Live Cells

**DOI:** 10.1101/2020.11.01.363663

**Authors:** Yutong Song, Peiyuan Huang, Xiaoying Liu, Bianxiao Cui, Liting Duan

## Abstract

Mitochondria, the powerhouse of the cell, are dynamic organelles that undergo constant morphological changes. Increasing evidence indicates that mitochondria morphologies and functions can be modulated by mechanical cues. However, the mechano-sensing and -responding properties of mitochondria and the correlation between mitochondrial morphologies and functions are unclear due to the lack of methods to precisely exert mechano-stimulation on and deform mitochondria inside live cells. Here we present an optogenetic approach that uses light to induce deformation of mitochondria by recruiting molecular motors to the outer mitochondrial membrane *via* light-activated protein-protein hetero-dimerization. Mechanical forces generated by motor proteins distort the outer membrane, during which the inner mitochondrial membrane can also be deformed. Moreover, this optical method can achieve subcellular spatial precision and be combined with other optical dimerizers and molecular motors. This method presents a novel mitochondria-specific mechano-stimulator for studying mitochondria mechanobiology and the interplay between mitochondria shapes and functions.

## Introduction

Mitochondria are double membrane-bound organelles known as the ‘powerhouse’ of eukaryotic cells that produce adenosine triphosphate (ATP). They also contribute to many other fundamental cellular processes including the synthesis of phospholipids and heme, reactive oxygen species (ROS) production, calcium homeostasis, cell growth and differentiation, cell cycle control, and cell death^1^. The constant dynamic changes of mitochondrial morphologies are influenced by delicate homeostasis between mitochondrial fission and fusion events by which mitochondria separate or merge. Mitochondria are also responsive to mechanical stimuli with their shapes and functions being affected and regulated. It has been observed that mechanical strains applied to the cells can lead to mitochondria elongation, the release of cytochrome c from mitochondria, decline of mitochondria membrane potential, and ROS production^2,3^. Moreover, external mechanical stresses are capable of directly and immediately initiating mitochondrial fission^3,4^. In addition, it has been found that mitochondria deformation happens during cardiac mechanical cycles^5,6^, exposure to osmotic pressure^7^, cell migration through narrow constrictions of microfabricated channels^8^, and localization within narrow axonal spaces in neurons^9^.

Despite mounting evidence indicating that mitochondria morphologies and functions can be modulated by mechanical forces, the mechano-sensing and -responding properties of mitochondria and the correlation between mitochondrial morphologies and functions are largely unclear. To answer these questions, a mechano-stimulating method is greatly desired that can apply force directly and precisely towards mitochondria with high organelle-specificity, non-invasiveness, good controllability, high-throughput, and compatibility with live-cell microscopy methods for simultaneous recording^8^. To date, very limited methods are available to mechanically perturb mitochondria. For instance, the atomic force microscope has been used to apply pressure to deform intracellular mitochondria through direct physical contact with the cell membrane using a cantilever^10^. Another method is bacteria-induced collisions which uses a motile bacteria, *Shigella flexneri*, to move throughout the host cells and randomly collide with and consequently deform intracellular mitochondria, however, in an uncontrollable and unpredictable manner^10^. Both methods suffer from low throughput, and other cellular components will be perturbed which may confound the final results.

The rapidly emerging optogenetic technologies provide new opportunities to establish targeted and non-invasive intracellular mechano-stimulation. Optogenetics combines optics and genetics to reversibly and precisely regulate protein functions in living cells with light. Different types of light-inducible protein-protein interactions, such as hetero-dimerization and homo-interaction, have been widely used to control biological activities including gene expression^11-14^ and signaling transduction^15-21^. In a very recent study, an optical tool based on a bacterial actin nucleation promoting factor has been developed to trigger actin polymerization and thus generate compressing force to deform mitochondria^22^. The mechanical force provided by molecular motors, a class of natural force-generating protein molecules that mobilize on the cytoskeleton, has been manipulated by light to drive the intracellular movement of organelles via light-inducible association between organelles and molecular motors^23-25^.

During our previous work that uses light to control the intracellular redistribution of organelles^24^, some stretching events of mitochondria were observed upon the light-induced association of molecular motors. Indeed, inside cells, the molecular motor can contribute to the change of mitochondria morphology. For example, molecular motors have been shown involved in the extension of mitochondria along microtubule arrays to facilitate sperm tail elongation^26^. Actually, the direct linkage between molecular motors and the lipid membrane shape has been reconstituted in artificial *in vitro* systems where synthesized Giant Unilamellar Vesicle (GUV) can be deformed by molecular motors which results in the formation of lipid tubes stretching out from GUV along microtubules^27^.

In this report, we set out to explore whether inside live cells, the mechanical forces generated by molecular motors are sufficient and efficient to deform mitochondrial membranes. Our results show that light-inducible intracellular mechano-stimulation and mitochondria deformation can be achieved by connecting the mitochondrial membrane to molecular motors with optical hetero-dimerizers, cryptochrome 2 (CRY2) and its binding partner CIBN. During the deformation of the outer mitochondrial membrane (OMM), the inner mitochondrial membrane (IMM) can also be deformed. Furthermore, optically induced mitochondria deformation can achieve spatial precision at the subcellular level. In addition, we show that this strategy is generally applicable to other optical hetero-dimerizer and different motors. We expect that our optical method will be a useful tool to facilitate the investigation of the interplay between mitochondria shapes and functions as well as the mechano-regulation of mitochondria.

## Results and Discussion

In our design, the photolyase homology region of CRY2 (a.a. 1-498) is fused to the transmembrane domain of mitochondrial Rho GTPase, Miro1, which targets the outer mitochondrial membrane^28^(Figure 1A). Kinesins are a type of motors that generate forces of several pico-newtons and can move toward the plus end of microtubules with a speed of several hundred nanometers per second^29^. The N-terminal region of CIBN, CIBN (a.a. 1-170) is fused to truncated kinesin-1 heavy chain isoform KIF5A which is unable to bind to cargos^30^. CRY2 and CIBN dimerize within milliseconds in the presence of blue light and dissociate in minutes after removal of blue light^31^. In our system, upon blue light stimulation, the motors are recruited to mitochondria via CRY2/CIBN association, thereby exerting mechanical forces provided by motors to deform mitochondria.

**Figure 1.**
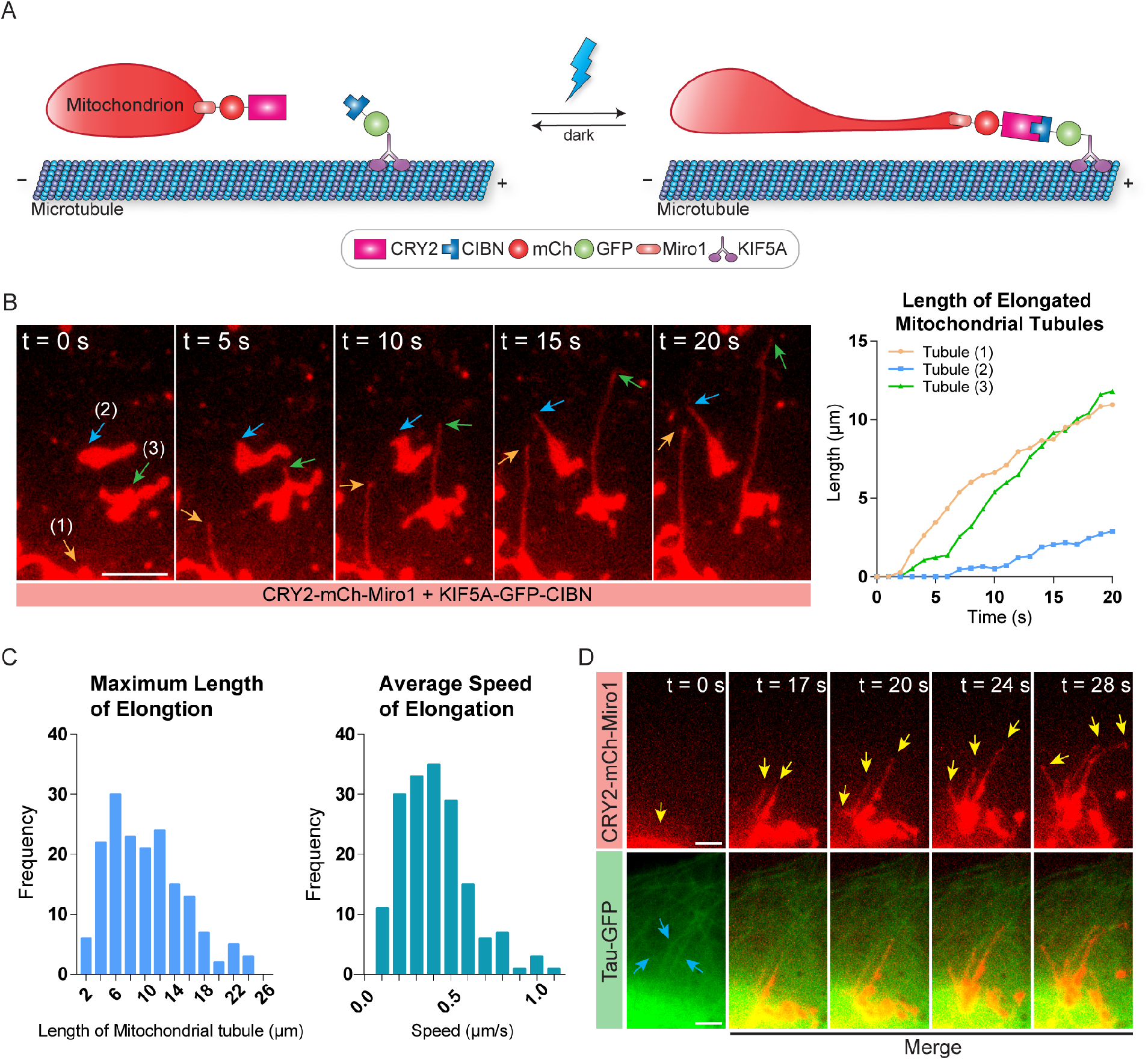
Optical induction of mitochondria deformation in living cells. (A) Schematic representation of light-controlled mitochondria deformation based on CRY2-CIBN heterodimerization. CRY2 is fused to mCh and Miro1, which anchors the fused protein to the outer mitochondrial membrane. CIBN is fused to GFP and KIF5A. Upon blue light exposure, CRY2 binds with CIBN, recruiting kinesins onto mitochondria. Kinesins mobilize toward the positive end of the microtubule and thus stretch the mitochondria. (B) Fluorescence images of mCh-tagged mitochondria showing deformation and elongation after blue light stimulation as indicated by arrows, in the COS-7 cell expressing CRY2-mCh-Miro1 and KIF5A-GFP-CIBN. The right graph maps the lengths of the elongating mitochondrial tubules with respect to time. (C) Quantification of the maximum length and the average speed of light-induced tubule elongation in a total of 171 individual mitochondria across 2 independent cultures that showed mitochondria deformation. (D) Mitochondria deform along underlying microtubules. The COS-7 cell was transfected with CRY2-mCh-Miro1, KIF5A-CIBN and TAU-GFP where TAU-GFP labels microtubules. Three mitochondrial tubules indicated by yellow arrows were extending after blue light stimulation along with the underlying microtubules marked by blue arrows. Scale bars, 5 μm.

### The deformation of mitochondria by optical recruitment of molecular motors

In COS-7 cells expressing CRY2-mCh-Miro1 and KIF5A-GFP-CIBN, upon blue light exposure, tubules could be seen stretched out from single mitochondria (Figure 1B, Movie S1). The lengths of the three extending tubules highlighted by arrows were measured, where the longest reached 17.8 μm while the shortest extended for 2.87 μm in 20 seconds. The whole-cell image of Figure 1B can be found in Figure S1A. It is worth noting that different cells may have varied extents of light-induced mitochondria deformation, and not all mitochondria with observed tubule formation deformed simultaneously after illumination. In some cells, we observed a good quantity of mitochondria undergoing significant membrane deformation during blue light exposure (Figure S1A), while in other cells only a portion of (Figure S1B), or very few (Figure S1C) mitochondria showed stretching. This variation can be possibly attributed to distinctive intracellular environments and differential physical properties of mitochondria in different cells. We quantified the stretching among cells with significant mitochondria deformation. 574 individual mitochondria among 12 cells from 6 independent experiments were measured. On average, 38.7% of mitochondria showed light-induced deformation within 120 seconds after blue light illumination. Deformed mitochondria showed discrepant lengths and speeds of light-induced tubule elongation. We also quantified the maximum length of extending tubules and the average speed of stretching in 171 mitochondria across 2 independent experiments. The average maximum length a tubule can reach is 10.1 μm. Surprisingly, the longest tubule can reach 25 μm, showcasing the great elasticity of mitochondria. The average speed of tubule elongation is 0.41 μm/sec, which agrees well with the movement speed of kinesin^29^ (Figure 1C).

To examine if the deformation of mitochondria is directed by microtubules, we labeled microtubules with GFP-tagged microtubule-associated protein Tau^32^. COS-7 cells were transfected with CRY2-mCh-Miro1, KIF5A-CIBN, and Tau-GFP. Upon light stimulation, merged images between mitochondrial tubules and microtubules showed that extending mitochondria tubules colocalized with underlying microtubule filament (Figure 1D), which confirmed that this light-induced deformation is dependent on the microtubule structure. In addition, we have demonstrated that light-inducible mitochondria deformation can be also applied to other types of cells, including HeLa, U2OS and 3T3 cells (Figure S2).

### Different phenomena of light-induced mitochondria stretching

In addition to stretching of a single mitochondrial tubule, we have also observed different types of deformation behaviors. This diversity is probably due to the complex arrangement of microtubule networks and the variable nature of kinesin-microtubule interaction. As shown in Figure 2A, some mitochondria had two tubules extending out from the main body in different directions. In another scenario as shown in Figure 2B, one stretching tubule diverged into two tubules and continued extending in different directions. To ask whether both scenarios result from the interactions between motors and the complex microtubule network, we examined the overlap between the elongating tubule and the microtubule alignment in COS-7 cells expressing CRY2-mCh-Miro1, KIF5A-CIBN, and Tau-GFP. As shown in Figure 2C, the two tubules extending from the same mitochondrion were indeed colocalized with the underlying microtubules, implying that two groups of kinesin motors on different separate domains of the same mitochondrial body interacted with two different microtubules and moved towards different directions.

**Figure 2.**
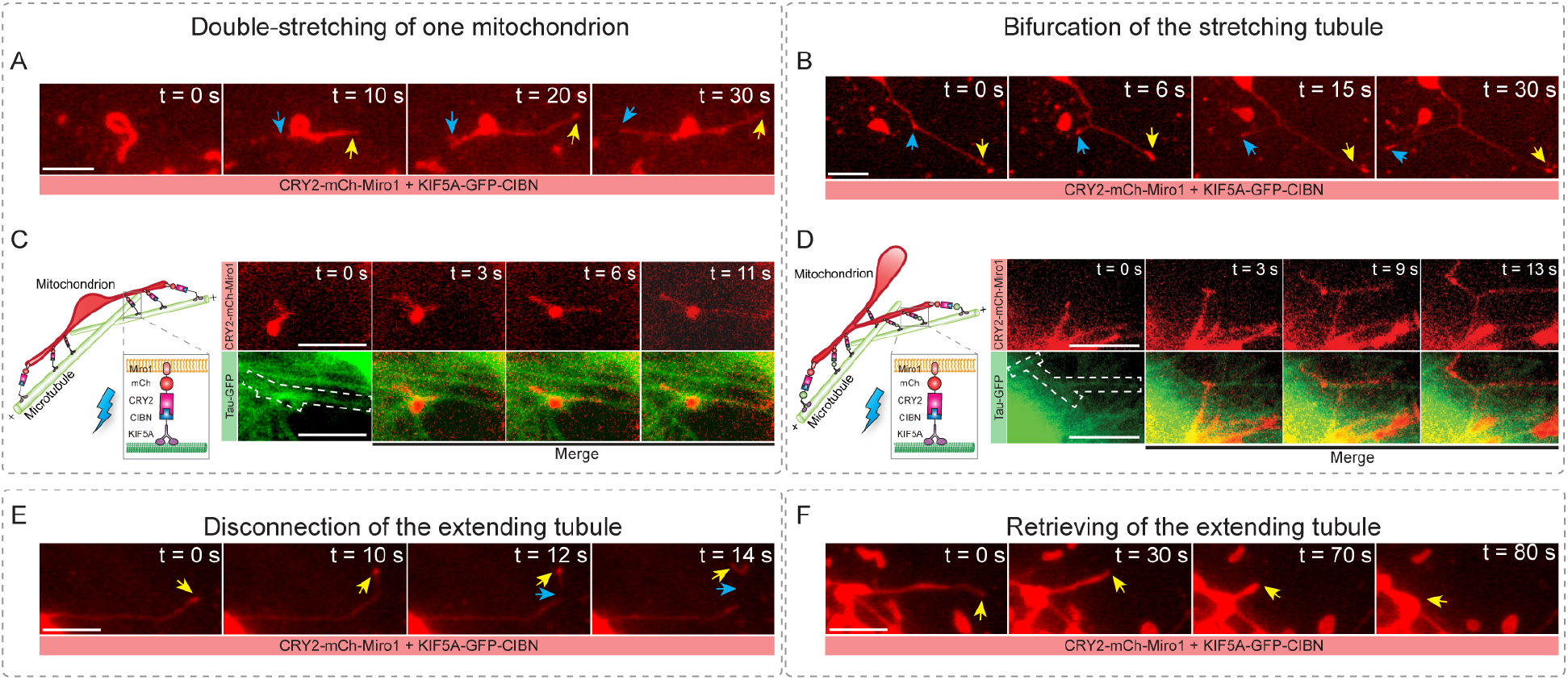
Different phenomena of light-induced mitochondria stretching. COS-7 cells were transfected with CRY2-mCh-Miro1 + KIF5A-GFP-CIBN (A,B,E,F), or CRY2-mCh-Miro1 + KIF5A-CIBN+TAU-GFP (C,D). (A) Two tubules indicated by arrows were stretching out in different directions from the same mitochondrial main body. (B) The extending mitochondrial tubule was split into two as indicated by arrows. (C) The colocalization of mitochondria and microtubules in merged images shows that the two tubules from the same mitochondrion were stretched in different directions along underlying microtubules highlighted by the dash-lined boxes. Schematic representation of this deformation is shown on the left. (D) Merged images showing a stretching mitochondrial tubule diverged at the crossover points between two microtubules indicated by the dashed-lined boxes. Schematic representation of this deformation is shown on the left. (E) The leading tip of an elongated tubule was disconnected away from the original mitochondrial body as indicated by arrows. (F) The tip of a stretched mitochondria tubule (marked by the arrow) was retracted back towards the main body. Scale bars, 5 μm.

Merged images of mitochondrial tubules and microtubules in Figure 2D shows that one extending tubule was split into two at the crossover of microtubule filaments, and the two new tubules continued to extend along the two microtubules in different directions. This bifurcation may be explained by the detachment of some kinesin molecules from the former microtubule and reattachment onto a new one at the crossover between two microtubule filaments, while remaining kinesins continued mobilizing on the original microtubule. This diverging extension agrees well with the results revealed in the simplified *in vitro* system in which motor-driven GUV tubules also showed bifurcation^27^. By quantifying 574 mitochondria with significant stretching within 120 sec intermittent blue stimulation among 12 cells across 6 independent experiments, our results showed that on average 8.2% of stretching mitochondria exhibited double-stretching while 15.8% had bifurcating tubules.

Upon continued light stimulation, the stretched mitochondria could often undergo a partition into two pieces or a retraction of the stretching tubule tip. As shown in figure 2E, when the tubule was highly elongated, the leading segment disconnected with the remaining mitochondrion, thus forming two individual parts. Following the disconnection, the break-away tip continued moving towards the periphery of the cell, while the mitochondria tubule usually stopped stretching and started retrieving because of the loss of driving force after losing the driving tip accumulated with motors. The retraction of the mitochondrial tubule allowed the stretching tubule to be shortened and retrieved back towards the mitochondrion body (Figure 2F). This retreat was probably due to loss of stability after the detachment flux of kinesin on the tubule surpassed the influx of kinesin caused by increased membrane tension during extension^27^. The retrieving phenomenon of some tubules also agrees with the results observed in GUV^27^. More examples for each of these five types of deformation are illustrated (Figure S3-S7).

### The deformation of the inner mitochondrial membrane during light-induced mitochondria stretching

IMM resides within the outer mitochondrial membrane (OMM) and encloses mitochondrial matrix. The mitochondrial matrix-targeting sequence, Mito^33^, was tagged by YFP to visualize the boundary of IMM (Figure 3A). OMM remained targeted by CRY2-mCh-Miro1, whereas KIF5A-CIBN contains no fluorescent label. In COS-7 cells expressing CRY2-mCh-Miro1, KIF5A-CIBN, and Mito-YFP, upon blue light exposure, while three long OMM tubules could be seen extending from different mitochondria, three IMM tubules from different mitochondria were stretching along the same path of their respective leading OMM tubule (Figure 3B, Movie S2). Lengths of OMM and IMM tubules were measured with respect to time, illustrating the simultaneous deformation of both membranes within the same mitochondrion. It is worth noting that OMM and IMM tubules can have different rates of elongation and disparate maximum lengths during the light-induced extension.

**Figure 3.**
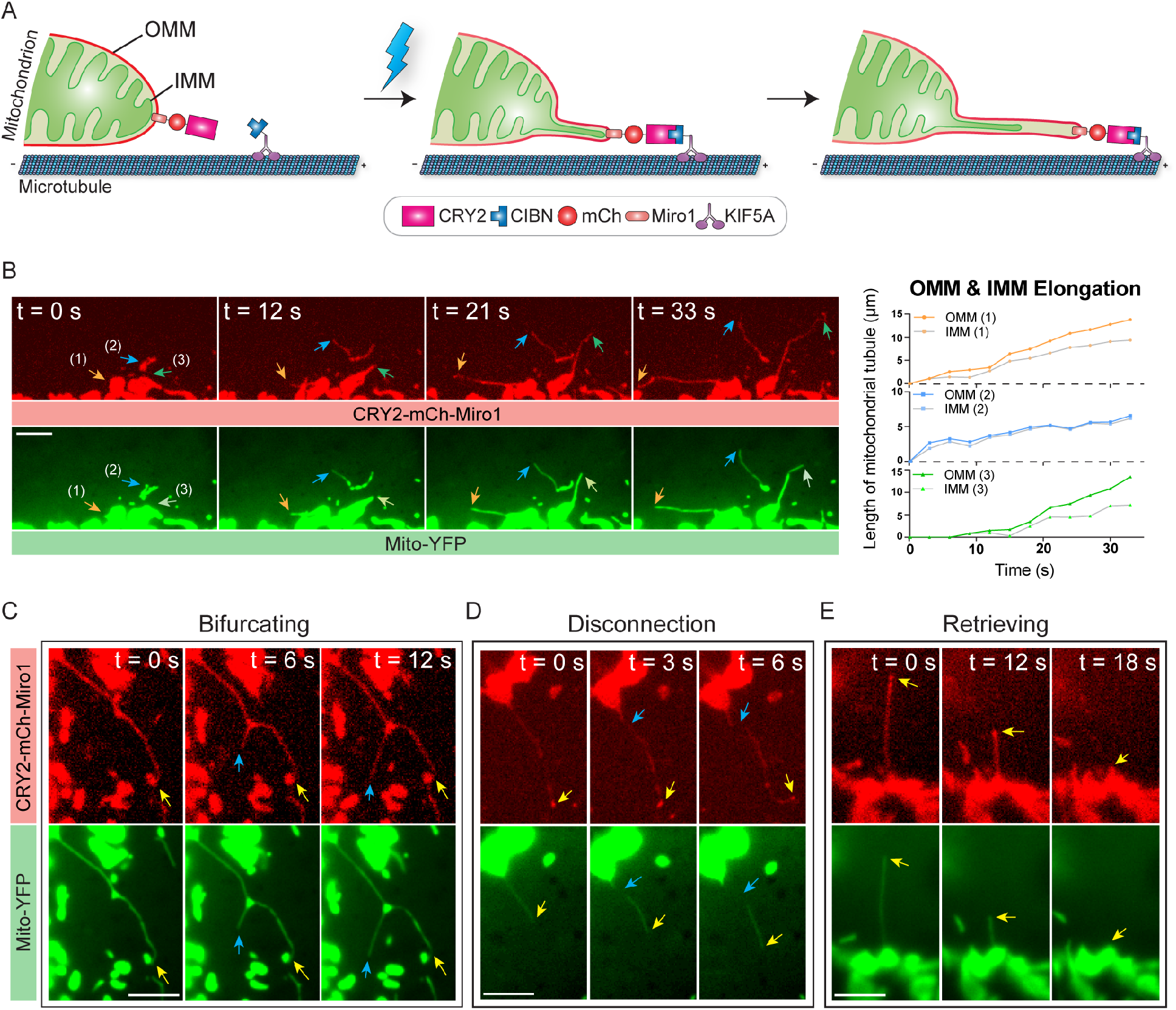
Deformation of inner mitochondrial membrane (IMM) during the light-induced outer mitochondrial membrane (OMM) reshaping. COS-7 cells were transfected with CRY2-mCh-Miro1, KIF5A-CIBN and Mito-YFP, where mCh signals localize on OMM and Mito-YFP targets the mitochondrial matrix to visualize IMM boundaries. (A) Schematic representation of IMM deformation following the light-induced stretching of OMM. (B) During light-induced stretching of OMM in the three mitochondria marked by arrows, the deformation and elongation of IMM tubules followed the same stretching paths indicated by arrows. The right graph quantifies the lengths of OMM and IMM tubules of the indicated three mitochondria with respect to time. (C) During the splitting of the extending OMM, IMM also diverged into two branches, as pointed by arrows. (D) Together with the disconnection of the OMM tubule from the mitochondrial body, the IMM tubule also got separated, as pointed by arrows. (E) While the tip of stretching OMM got retrieved, IMM was retracted as well as marked by arrows. Scale bars, 5 μm.

Interestingly, the lag in IMM deformation relative to OMM could be very significant. As shown in Figure S8, the OMM tubule increased from 1.6 to 19.8 μm after 57 s of intermittent blue light exposure, while the length of the IMM tubule maximized at 9.6 μm. On the contrary, in some cases, it was found that the IMM showed almost no deformation when its OMM had significant tubule extension (Figure S9). Quantification of 231 mitochondria among 11 cells across 2 independent experiments showed that on average 25.8% of mitochondria with OMM deformation did not show concurrent deformation of IMM. Such a separation between OMM and IMM, which is also reported in mitochondria residing inside narrow axon sections in neurons^9^, suggests different physical properties of different membranes which need further investigation.

Like OMM, IMM can undergo bifurcating, retrieving, and disconnection (Figure 3C-E) together with the corresponding OMM. More examples for each type of deformation behaviors are illustrated in Figure S9-S13. Intriguingly, during the partition of stretching mitochondria, the Mito-YFP remained inside the inner membrane space, indicating that the structure of IMM remained intact without any physical ruptures. The nature of the partition induced in these deformed mitochondria is worth further inspection.

### The spatial control of light-inducible mitochondria deformation

One exceptional advantage of optical control is the subcellular spatial resolution. We demonstrated that the deformation of mitochondrial membranes could be restricted to a specific subcellular area (Figure 4). In the COS-7 cell expressing CRY2-mCh-Miro1 and KIF5A-GFP-CIBN, blue light was given to only the area indicated by the blue rectangle. Shapes of mitochondria that were not exposed to blue light in the dashed yellow rectangle remained in the original states. Inside the area illuminated by blue light, significant mitochondria stretching and elongation could be observed. Zoom-in images showed the difference between these two regions where only mitochondria within the illuminated area underwent deformation. Subsequent blue light illumination on the entire cell showed that tubule formation also occurred in mitochondria which were not stimulated in the first round of light stimulation, confirming that precise induction of membrane deformation in specific mitochondria was the result of spatial control.

**Figure 4.**
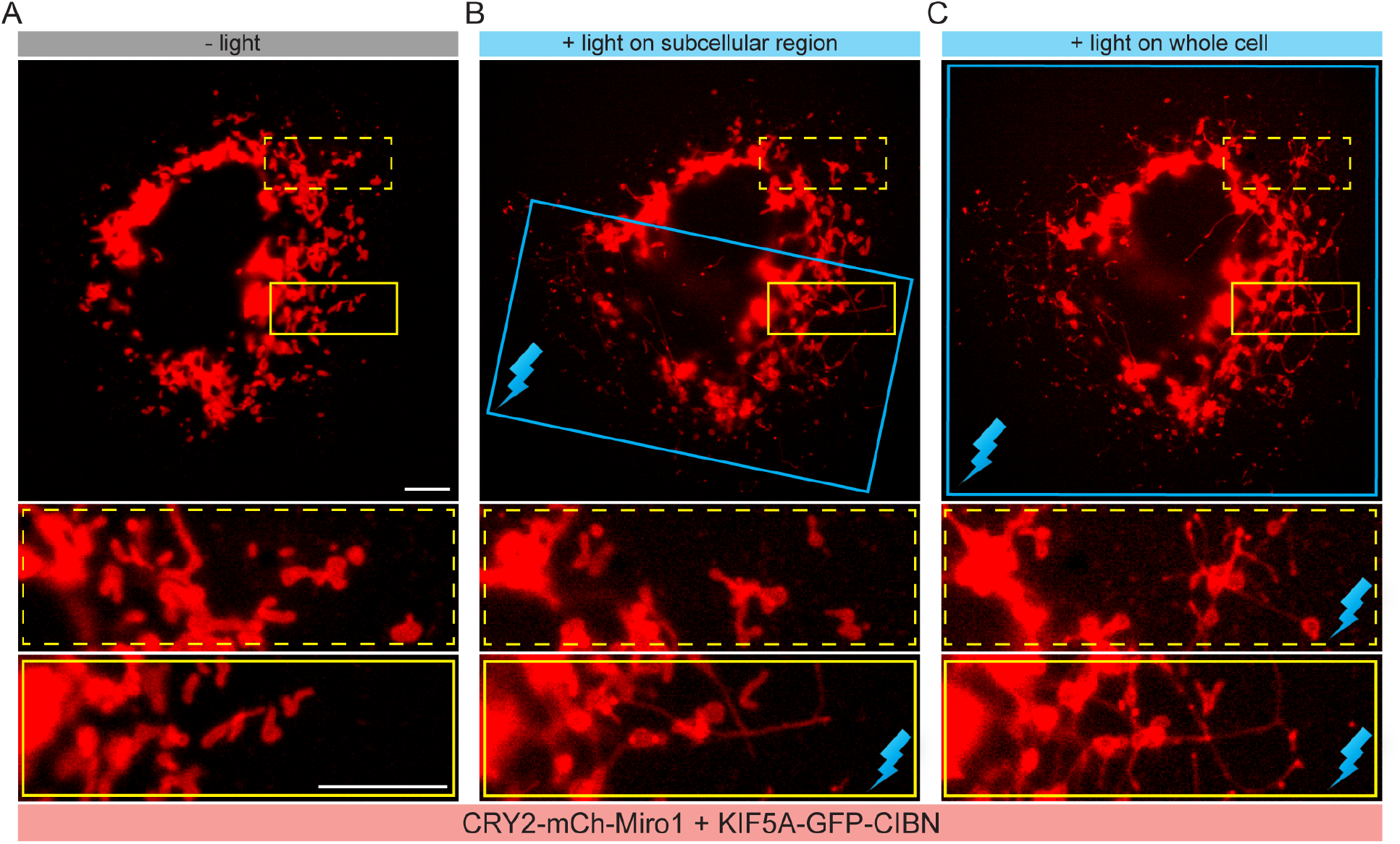
Spatial control of mitochondria membrane deformation in subcellular areas. The COS-7 cell was transfected with CRY2-mCh-Miro1 and KIF5A-GFP-CIBN. (A) Before any blue light stimulation, mitochondria stayed in short and chubby shapes, as shown in zoomed-in images of both dashed and solid yellow line-indicated areas. (B) Blue light was given to a subcellular region which is indicated by the blue box and contains the solid line indicated area. Tubule formation can only be observed in mitochondria within the illuminated area, as observed in the zoomed-in image of the solid line indicated area. (C) Subsequent illumination on the whole cell induced mitochondria deformation across the entire cell, as demonstrated in zoomed-in figures of both dashed and solid yellow line-indicated areas. Scale bars, 10 μm.

### The compatibility of the light-inducible mitochondria system with other optical dimerizers and molecular motors

After confirming that optical recruitment of kinesin 1 onto organelles is able to induce mitochondria deformation, we ask whether this design can be applied with other optical hetero-dimerizers and motors. In addition to CRY2-CIBN, there are other optical hetero-dimerizing pairs such as the improved light-induced dimer iLID and its binding partner SspB which are derived from the *Avena sativa* light-oxygen-voltage 2 (AsLOV2) domain^34^. To integrate iLID-SSpB into our design, we first fused the iLID to Miro1 and SspB(Micro), a version of SspB with medium binding affinity, to KIF5A (Figure 5A). In COS-7 cells expressing iLID-mCh-Miro1 and KIF5A-GFP-SspB(Micro), tubule deformation could be successfully induced in mitochondria when activated by blue light (Figure 5B). Then we exchanged the position of the dimerizing proteins, connecting SspB(Micro) to Miro1 and iLID to KIF5A (Figure S14A). Results showed that tubules could still be stretched out from mitochondria upon blue light exposure (Figure S14B), suggesting that altering positions of the iLID/SspB pair did not affect its ability to induce deformation. Additionally, we confirmed that another version of SspB, SspB(Nano), which exhibits a varied binding ability to iLID protein, worked as well in our system in prompting mitochondrial tubule formation (Figure S14C).

**Figure 5.**
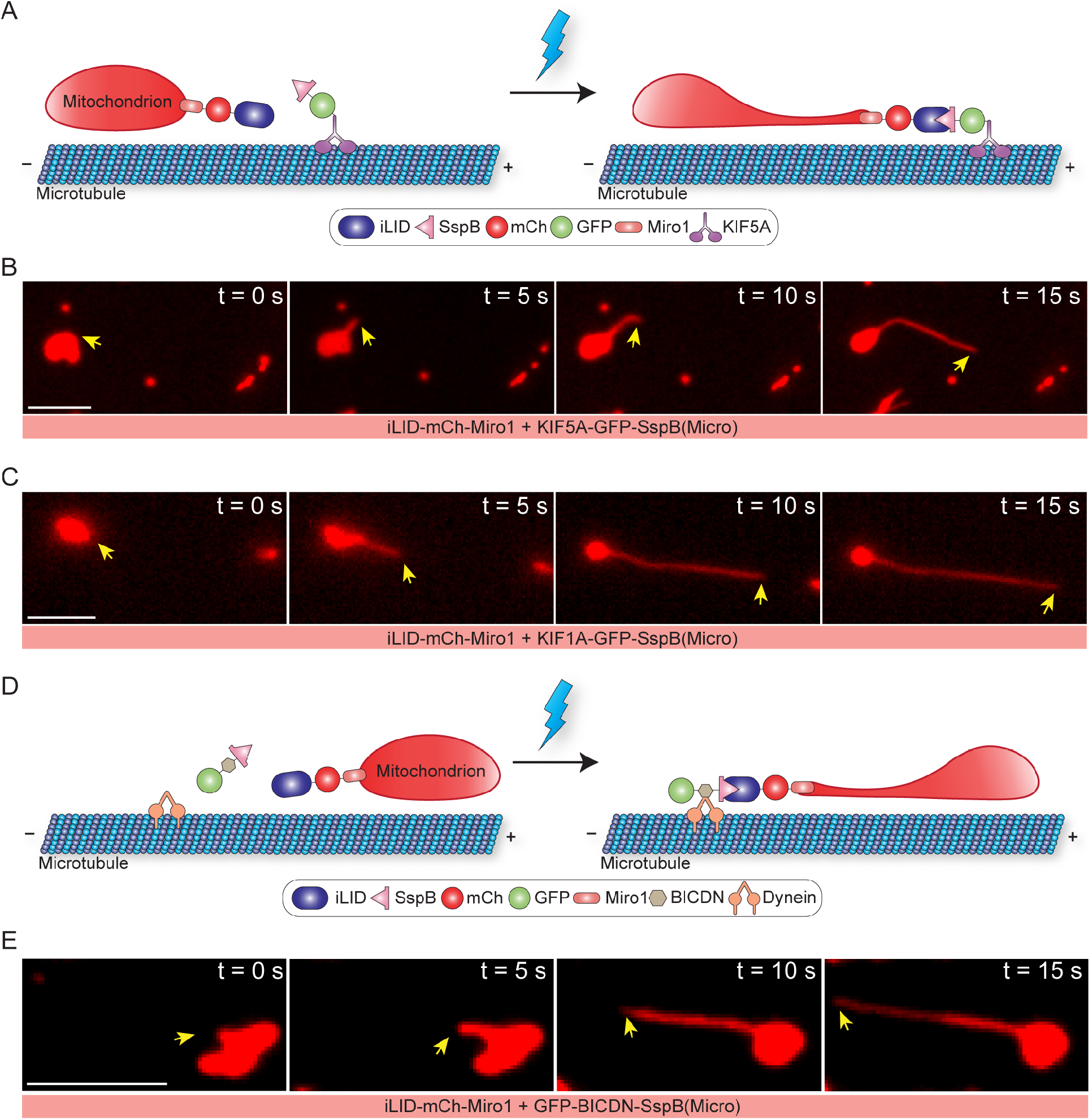
The light-inducible membrane deformation system is generally applicable to other optical dimerizers and molecular motors. (A) Scheme for light-induced mitochondria deformation using the iLID-SspB module. In this design, iLID is targeted to the OMM by Miro1 and SspB(Micro) is fused to kinesin. (B) After blue light stimulation, mitochondria exhibited stretching deformation in the COS-7 cell expressing iLId-mCh-Miro1 and KIF5A-GFP-SspB(Micro). (C) By integrating KIF1A, the truncated isoform of kinesin-3 into the iLID-SspB based system, optical recruitment of kinesin-3 can also induce mitochondria deformation in the COS-7 cell expressing iLID-mCh-Miro and KIF1A-GFP-SspB(Micro). (D) Scheme for optical induction of mitochondria deformation using the mechanical force provided by dynein motors that mobilize toward the minus end of microtubules. In this design, SspB(Micro) is fused to BICDN which is a dynein adapter protein. (E) The dynein-driven optogenetic system can induce mitochondria deformation upon blue light stimulation in the COS-7 cell expressing iLID-mCh-Miro1 and GFP-BICDN-SspB(Micro). Scale bars, 5 μm.

We also implemented different types of motors in the light-induced mitochondria deformation system. Both kinesin 1 and kinesin 3 belong to the kinesin family that move on microtubules towards the plus end. We thus chose to fuse the truncated isoform of kinesin 3, KIF1A, to SspB(Micro). Fluorescence images showed that light-induced recruitment of KIF1A was also able to induce mitochondria deformation in the COS-7 cell expressing iLid-mCh-Miro1 and KIF1A-GFP-SspB(Micro) (Figure 5C). To test whether another type of microtubule-associated motor protein, the minus-end-directed dynein can drive the organelle deformation, we fused the dynein-interacting sequence, BICDN, to SspB(Micro) to construct a system that can optically link the mitochondrial membrane with dynein (Figure 5D). After blue light stimulation, OMM showed tubule elongation similar to what was elicited by kinesin (Figure 5E). All these results demonstrate the good generalizability and applicability of our strategy to optically induce mitochondria membrane deformation by combining with different optogenetic toolkits and different motors.

## Discussion

In this report, we demonstrate that mitochondria membrane deformation can be optically and non-invasively induced in live cells via light-induced recruitment of molecular motors onto mitochondria. We show that both the outer and inner membranes of mitochondria can be stretched and deformed using this optical strategy. Different phenomena following the initial deformation of inner and outer membranes are observed, including extension of several tubules from the same mitochondria, the bifurcation of the stretching tubule, retrieval of the stretching tip, and the partition of the stretching fragment. Subcellular spatial control can be achieved by confining the light stimulation area. In addition, we also show that different optical hetero-dimerizers and molecular motors can be incorporated using this strategy to induce mitochondria distortion.

We have demonstrated that different optogenetic modules, CRY2/CIBN and iLID/SspB, can be implemented into our system to trigger mitochondria deformation. The results indicate that our optogenetic control system is open for integration of other optical dimerizers. For example, optical hetero-dimerizers of different kinetics and binding affinities can be chosen to construct customized systems for inducible mitochondria deformation. In addition, optical hetero-dimerizers activatable by different wavelengths of light, such as PHYB-PIF6 and Bphp-PpsR2 that is sensitive to red light or UVR8-COP1 that can be stimulated by UV light, can be used to satisfy distinct illumination conditions. It is also possible to construct an orthogonal platform to independently modulate the deformation of mitochondria and other cellular processes, by choosing hetero-dimerizers with well separated excitation spectra. Therefore, the adaptability of this system grants more possibilities of selecting different optogenetic modules for specific purposes according to various experimental needs. The flexibility of components in this optical control system also applies to motor molecules. Our report has proven that both kinesin and dynein can induce membrane deformation. Thus, we expect that other motors and motor isoforms may be integrated in this system to have similar effects as well, such as members of the myosin family. Expansion of the motor pool could assist future development of new control systems to tackle questions under different cellular conditions.

There are certain limitations in our system of light-inducible mitochondrial deformation. Firstly, it is hard to control the exact location and direction of induced membrane deformation due to the critical dependence on the underlying microtubule structures. For light-induced deformation to occur, microtubules are required to be present at the sites of the mitochondria to direct and support motor movement. As a result, whether a specific organelle membrane or membrane region can be reshaped and the stretching direction largely depend on the naturally formed microtubule structures which vary from cell to cell, and location to location and cannot be artificially sculpted by any method available up to date. In addition, our method is not effective in inducing the deformation of every mitochondria. Some mitochondria that established light-induced interaction with microtubules showed movement of the entire mitochondrial body instead of obvious deformation. The factors that decide whether and how much a mitochondrion will be stretched by optical recruitment of molecular motors need further investigation. Furthermore, limitation also exists in the quantification of actual association between membrane and motors. It is hard to quantitatively assess or precisely control the number of molecular motors recruited to the membrane after light activation. Thus, the total force exerted by motors on the membrane is difficult to quantify.

In conclusion, we demonstrate an optical method that can remotely and non-invasively deform mitochondria inside living cells, with minimal perturbation towards other cellular components. We expect this method can greatly facilitate our understanding on how mitochondria sense and respond to mechanical forces, the physical attributes of mitochondrial membranes, and the functional significance of mitochondrial morphologies.

## Methods

### Plasmids Construction

All the plasmids used in this study were cloned in the mammalian expression vector pEGFPN1 or pmCherryC1. CRY2-mCh-Miro1 and KIF5A-GFP-CIBN were from previous work^24^. iLID-mCh-Miro1 was made by inserting iLID to and replacing CIBN in CIBN-mCh-Miro1 using In-Fusion (Clonetech). SspB(Micro)-mCh-Miro1 and SspB(Nano)-mCh-Miro1 were made in a similar way, by replacing CIBN in CIBN-mCh-Miro1 with SspB(Micro) or SspB(Nano) respectively using In-Fusion. KIF5A-GFP-SspB(Micro) and KIF5A-GFP-iLID were constructed by inserting SspB(Micro) and iLID into KIF5A-GFP-CIBN respectively to replace CIBN using In-Fusion. Motor domain of rat KIF1A (a.a 1-383) was inserted to KIF5A-GFP-SspB(Micro) and replaced KIF5A to make KIF1A-GFP-SspB(Micro) using InFusion. GFP-BICDN-SspB(Micro) was constructed by inserting SspB(Micro) to GFP-BICDN using In-Fusion. Tau-GFP was constructed by replacing YFP in Tau-YFP with GFP using ligation. The details for plasmid constructions are summarized in Table S1.

### Cell culture and transfection

COS-7 (ATCC® CRL-1651™) cells were cultured in DMEM medium (Thermo Fisher Scientific) supplemented with 10% FBS (fetal bovine serum, Clontech) and 1% P/S (Penicillin-Streptomycin, Thermo Fisher Scientific). All cell cultures were maintained at 37°C with 5% CO_2_. Cells were plated on 35 mm confocal dishes with hole size of 13Ø or 20Ø (SPL) and allowed to grow 1-2 days before transfection. All cells were transfected with desired DNA plasmids using Lipofectamine 2000 (Thermo Fisher Scientific) according to the manufacturer’s protocol. Transfected cells were allowed to recover and express the desired proteins overnight in a complete culture medium. Fluorescence imaging of the transfected cells was conducted a day after transfection. HeLa, U2OS and 3T3 cells were cultured and transfected following the same protocol as that for COS-7 cells described above.

### Fluorescence Imaging

All live imaging was performed on an epifluorescence microscope (Leica DMi8S, Thunder Imager) with an on-stage CO_2_ incubator and a motorized stage. An adaptive focus control was used during the whole imaging process to keep the region of interest in focus. Imaging experiments were conducted one day after cell transfection. For blue-light stimulation, pulsed blue light (200 ms pulse duration at 1-s or 5-s intervals at 9.7 W/cm2) was used for GFP imaging and to initiate CRY2-CIB1 interactions. For mCh imaging, pulsed green light (200 ms pulse duration) was used. The light intensity was measured by THORLABS PM100D optical power and energy meter right above the 100x objective.

### Data analysis

Measurement of the maximal lengths and extending speeds of mitochondrial tubules was done on raw fluorescence images using ImageJ. We measured the length of stretched mitochondrial membrane tubules and the average speed of elongation of 171 individual mitochondria that showed significant deformation from 3 independent experiments. All selected mitochondria and their tubules could be individually identified and tracked throughout the time lapse of the measurement. We defined the length of the tubule (in microns) as the distance between the leading tip and the point where the tubule stemmed from the mitochondrial main body along the path of the tubule. The maximum length of the tubule was defined as the longest a tubule could reach before halting, disconnection or retraction. Tubule lengths were measured on single frames. Average speed of tubule elongation was calculated as the quotient of maximum length and time interval (in seconds). All measurements of the deformation of both outer and inner mitochondrial membranes presented in the results were conducted following this method.

Data of mitochondria deformation statistics were acquired by counting and measuring mitochondria that could be individually identified and were separated from mitochondrial clusters in raw fluorescence images using ImageJ. For the analysis of different phenomena of light-induced mitochondria stretching, we chose 12 sample cells from 6 independent experiments and counted 574 individually-identifiable mitochondria in total. Among selected mitochondria in each cell, we counted and recorded the number of mitochondria that showed significant deformation in the first 2 minutes after blue light activation. We defined significant deformation as having a stretching tubule with an elongation of at least one time or more the length of the diameter of the mitochondrial main body. In addition to counting mitochondria with deformation in general, we also recorded the number of events of different stretching phenomena in the same data set. For each cell, we calculated the percentage of mitochondria that showed significant deformation, as well as the occurring frequency of each stretching phenomenon we described. Total average of the results from the 12 cells was reported.

The analytical process for the statistics of IMM deformation during light-induced mitochondria stretching was similar, with the difference being only mitochondria with significant deformation of outer membrane were selected. 11 sample cells from 2 independent experiments were chosen and 231 deformed mitochondria were counted in total. Among selected mitochondria in each cell, we counted and recorded the number of mitochondria that showed each of the four types of inner membrane phenomena in the first 2 minutes after blue light activation. Calculation of the occurring frequency of each type of inner mitochondrial membrane deforming phenomenon was done by Microsoft Excel. Total average of the results from the 11 cells was reported.

## Acknowledgements

We thank Dr. Chandra Tucker (University of Colorado Denver) for providing backbone plasmids for CIB1 and CRY2 constructs; we thank Dr. Xinnan Wang (Stanford University) for providing Miro1 plasmid; we thank Dr. Casper Hoogenraad (Utrecht University) for providing plasmids encoding the truncated motor construct KIF5A, dynein/dynactin adapter BICDN. This work was supported by a Direct Grant from the Chinese University of Hong Kong (4055095), Shun Hing Institute of Advanced Engineering (SHIAE) Grant (4720247) and a General Research Fund (GRF)/Early Career Scheme (ECS) (24201919) from the Research Grants Council (RGC) in Hong Kong.

## Author Contributions

L.D., B.C. and Y.S. conceived the project and designed the experiments. P.H., X.L. and Y.S. conducted the experiments. Y.S. and X.L. analyzed the data. Y.S. and L.D. wrote the paper. All authors contributed to the discussion of the results.

## References

1. McBride, H. M., Neuspiel, M. & Wasiak, S. Mitochondria: More Than Just a Powerhouse. Current Biology vol. 16 R551–R560 (2006).

2. Chapman, K. E. et al. Cyclic mechanical strain increases reactive oxygen species production in pulmonary epithelial cells. Am. J. Physiol. Lung Cell. Mol. Physiol. 289, L834–41 (2005).

3. Bartolák-Suki, E., Imsirovic, J., Nishibori, Y., Krishnan, R. & Suki, B. Regulation of Mitochondrial Structure and Dynamics by the Cytoskeleton and Mechanical Factors. Int. J. Mol. Sci. 18, (2017).

4. Feng, Q. & Kornmann, B. Mechanical forces on cellular organelles. Journal of Cell Science vol. 131 jcs218479 (2018).

5. Rog-Zielinska, E. A., O’Toole, E. T., Hoenger, A. & Kohl, P. Mitochondrial Deformation During the Cardiac Mechanical Cycle. Anat. Rec. 302, 146–152 (2019).

6. Yaniv, Y. et al. Analysis of mitochondrial 3D-deformation in cardiomyocytes during active contraction reveals passive structural anisotropy of orthogonal short axes. PLoS One 6, e21985 (2011).

7. Javadov, S., Chapa-Dubocq, X. & Makarov, V. Different approaches to modeling analysis of mitochondrial swelling. Mitochondrion 38, 58–70 (2018).

8. Feng, Q., Lee, S. S. & Kornmann, B. A Toolbox for Organelle Mechanobiology Research— Current Needs and Challenges. Micromachines vol. 10 538 (2019).

9. Fischer, T. D., Dash, P. K., Liu, J. & Waxham, M. N. Morphology of mitochondria in spatially restricted axons revealed by cryo-electron tomography. PLoS Biol. 16, e2006169 (2018).

10. Helle, S. C. J. et al. Mechanical force induces mitochondrial fission. Elife 6, (2017).

11. Liu, H., Gomez, G., Lin, S., Lin, S. & Lin, C. Optogenetic control of transcription in zebrafish. PLoS One 7, e50738 (2012).

12. Polstein, L. R. & Gersbach, C. A. A light-inducible CRISPR-Cas9 system for control of endogenous gene activation. Nat. Chem. Biol. 11, 198–200 (2015).

13. Taslimi, A. et al. Optimized second-generation CRY2-CIB dimerizers and photoactivatable Cre recombinase. Nature Chemical Biology vol. 12 425–430 (2016).

14. Wang, X., Chen, X. & Yang, Y. Spatiotemporal control of gene expression by a light-switchable transgene system. Nat. Methods 9, 266–269 (2012).

15. Zhang, K. et al. Light-mediated kinetic control reveals the temporal effect of the Raf/MEK/ERK pathway in PC12 cell neurite outgrowth. PLoS One 9, e92917 (2014).

16. Duan, L. et al. Optical Activation of TrkA Signaling. ACS Synth. Biol. 7, 1685–1693 (2018).

17. Grusch, M. et al. Spatio-temporally precise activation of engineered receptor tyrosine kinases by light. The EMBO Journal vol. 33 1713–1726 (2014).

18. Chang, K.-Y. et al. Light-inducible receptor tyrosine kinases that regulate neurotrophin signalling. Nature Communications vol. 5 (2014).

19. Toettcher, J. E., Weiner, O. D. & Lim, W. A. Using optogenetics to interrogate the dynamic control of signal transmission by the Ras/Erk module. Cell 155, 1422–1434 (2013).

20. Ong, Q. et al. The Timing of Raf/ERK and AKT Activation in Protecting PC12 Cells against Oxidative Stress. PLoS One 11, e0153487 (2016).

21. Huang, P. et al. Optical Activation of TrkB Signaling. J. Mol. Biol. 432, 3761–3770 (2020).

22. Spier, A. et al. Bacterial FtsZ induces mitochondrial fission in human cells. doi:10.1101/2020.01.24.917146.

23. van Bergeijk, P., Adrian, M., Hoogenraad, C. C. & Kapitein, L. C. Optogenetic control of organelle transport and positioning. Nature 518, 111–114 (2015).

24. Duan, L. et al. Optogenetic control of molecular motors and organelle distributions in cells. Chem. Biol. 22, 671–682 (2015).

25. Harterink, M. et al. Light-controlled intracellular transport in Caenorhabditis elegans. Curr. Biol. 26, R153–4 (2016).

26. Noguchi, T., Koizumi, M. & Hayashi, S. Sustained elongation of sperm tail promoted by local remodeling of giant mitochondria in Drosophila. Curr. Biol. 21, 805–814 (2011).

27. Leduc, C., Campàs, O., Joanny, J.-F., Prost, J. & Bassereau, P. Mechanism of membrane nanotube formation by molecular motors. Biochim. Biophys. Acta 1798, 1418–1426 (2010).

28. Fransson, S., Ruusala, A. & Aspenström, P. The atypical Rho GTPases Miro-1 and Miro-2 have essential roles in mitochondrial trafficking. Biochem. Biophys. Res. Commun. 344, 500–510 (2006).

29. Mallik, R. & Gross, S. P. Molecular motors: strategies to get along. Curr. Biol. 14, R971–82 (2004).

30. Cai, D., Hoppe, A. D., Swanson, J. A. & Verhey, K. J. Kinesin-1 structural organization and conformational changes revealed by FRET stoichiometry in live cells. J. Cell Biol. 176, 51–63 (2007).

31. Kennedy, M. J. et al. Rapid blue-light-mediated induction of protein interactions in living cells. Nat. Methods 7, 973–975 (2010).

32. Hoover, B. R. et al. Tau mislocalization to dendritic spines mediates synaptic dysfunction independently of neurodegeneration. Neuron 68, 1067–1081 (2010).

33. Rizzuto, R., Brini, M., Pizzo, P., Murgia, M. & Pozzan, T. Chimeric green fluorescent protein as a tool for visualizing subcellular organelles in living cells. Curr. Biol. 5, 635–642 (1995).

34. Guntas, G. et al. Engineering an improved light-induced dimer (iLID) for controlling the localization and activity of signaling proteins. Proc. Natl. Acad. Sci. U. S. A. 112, 112–117 (2015).

