## Supplemental materials for "Light-inducible Deformation of Mitochondria in Live Cells"

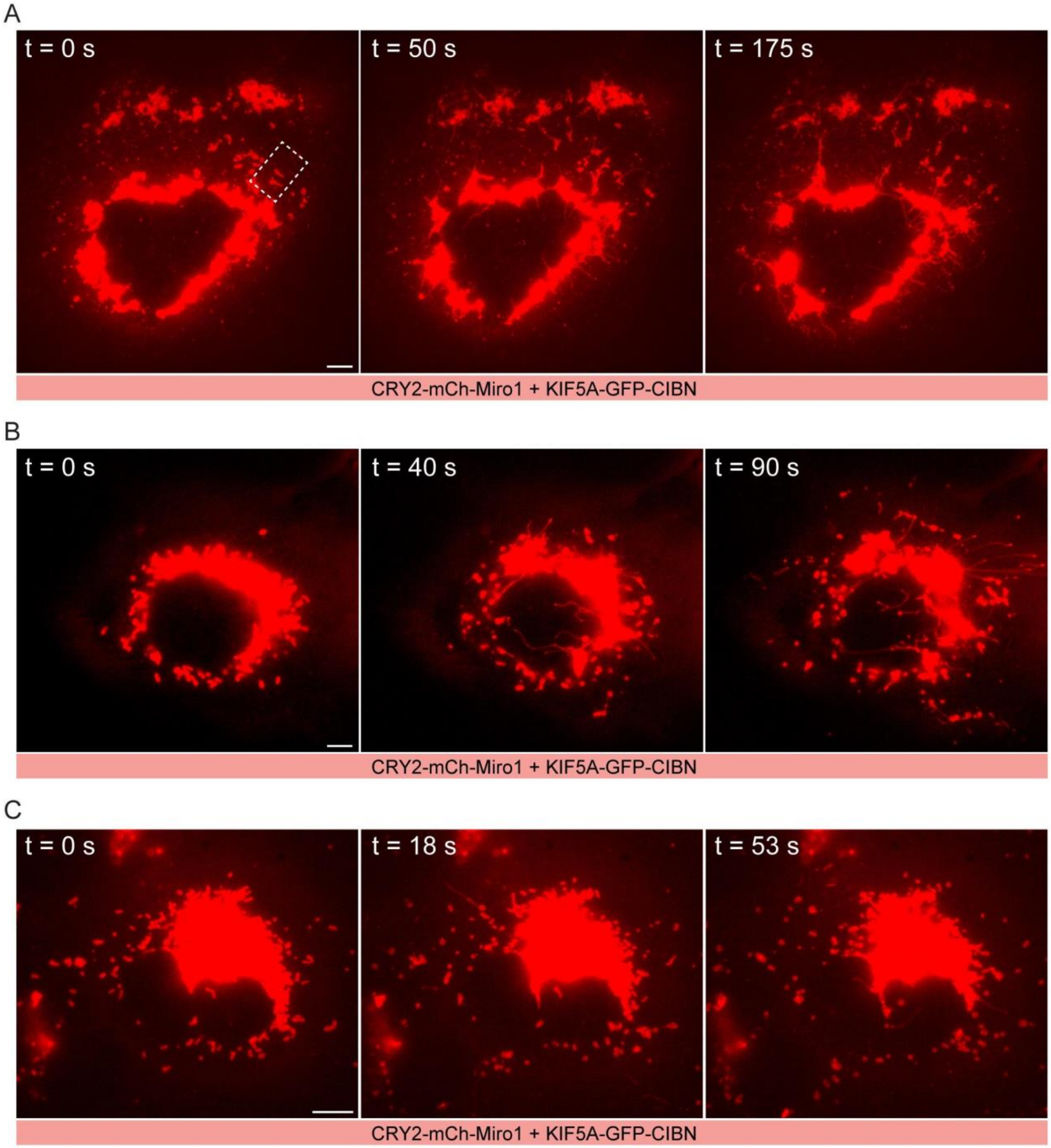


Figure S1. At the whole-cell level, some cells have many events of light-induced mitochondria deformation (A), some have multiple (B), while some have only a few (C). The region indicated by the dash-lined box in (A) is the same region shown in Figure 1B. The COS-7 cell was transfected with CRY2-mCh-Miro1 and KIF5A-GFP-CIBN. Scale bars, 5 µm.

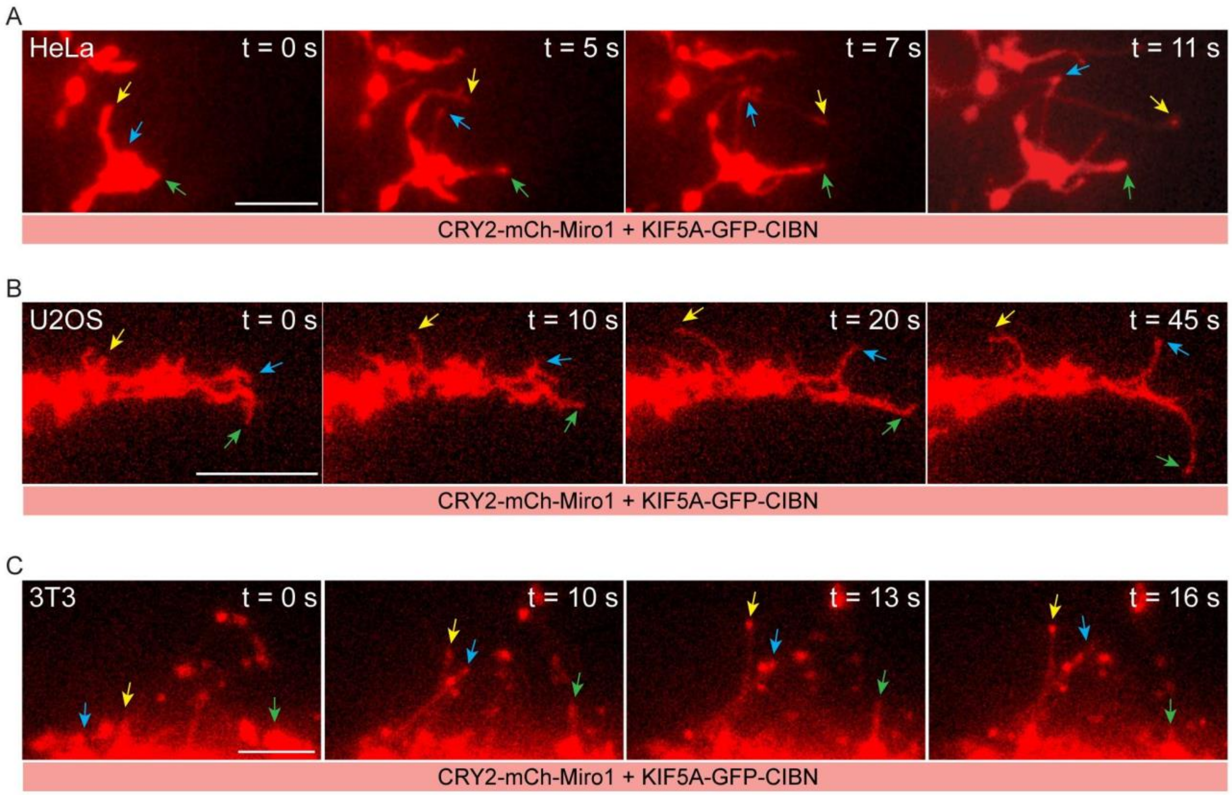


Figure S2. Light-inducible mitochondria deformation can be applied to different types of cell lines. (A) Deformation of mitochondria in HeLa cells upon blue light exposure. (B) Deformation of mitochondria in U2OS cells upon blue light exposure. (C) Deformation of mitochondria in 3T3 cells upon blue light exposure. All cells are transfected with CRY2-mCh-Miro1 and KIF5A-GFP-CIBN. Scale bars, 5 μm.

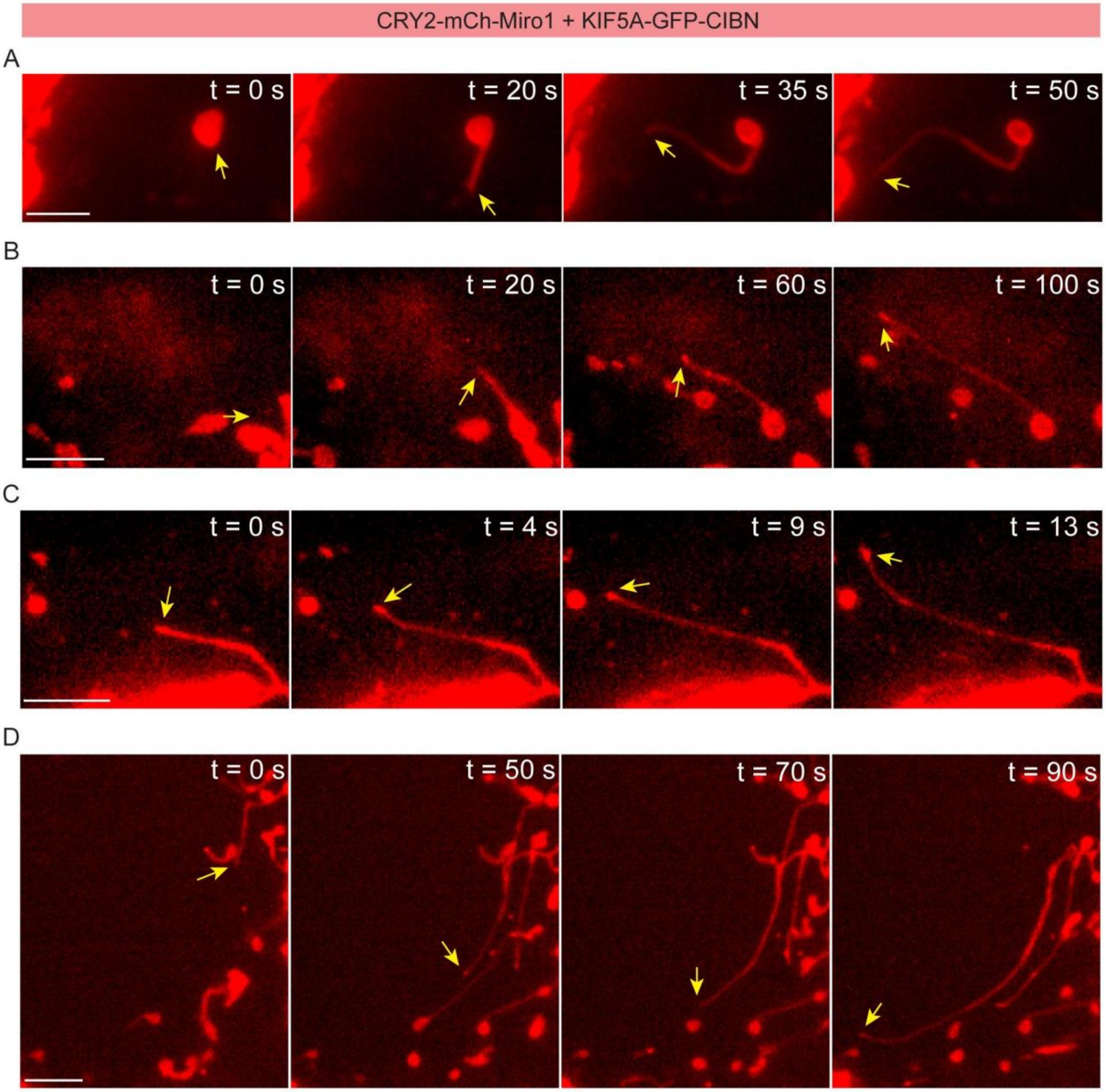


Figure S3. (A-D) Additional examples of deformation of mitochondria that show a single tubule stretching after blue light activation. Scale bars, 5 μm.

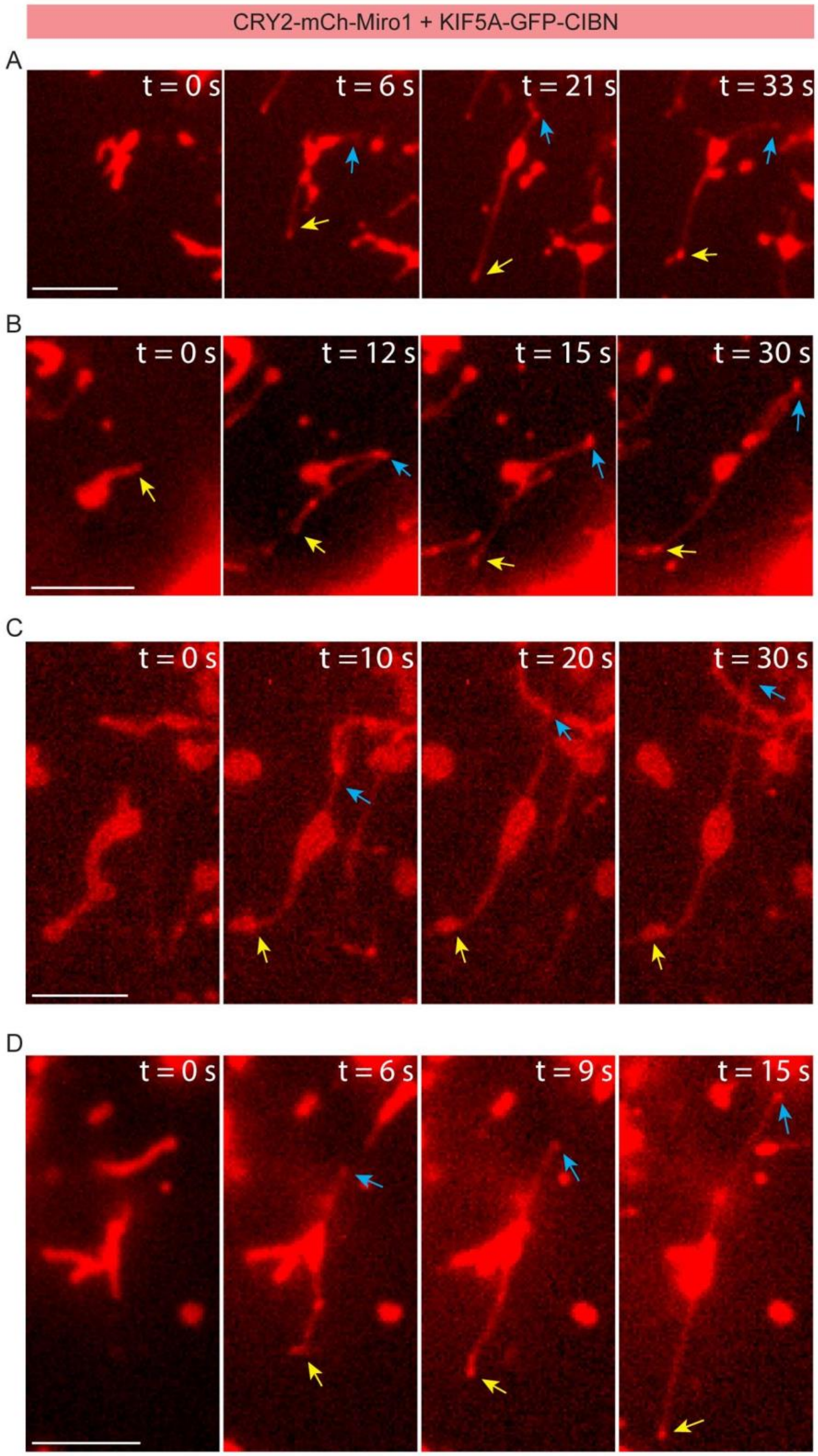


Figure S4. (A-D) Additional examples of mitochondrial outer membranes that show double stretching of tubules from one mitochondrion after blue light activation. Scale bars, 5 μm.

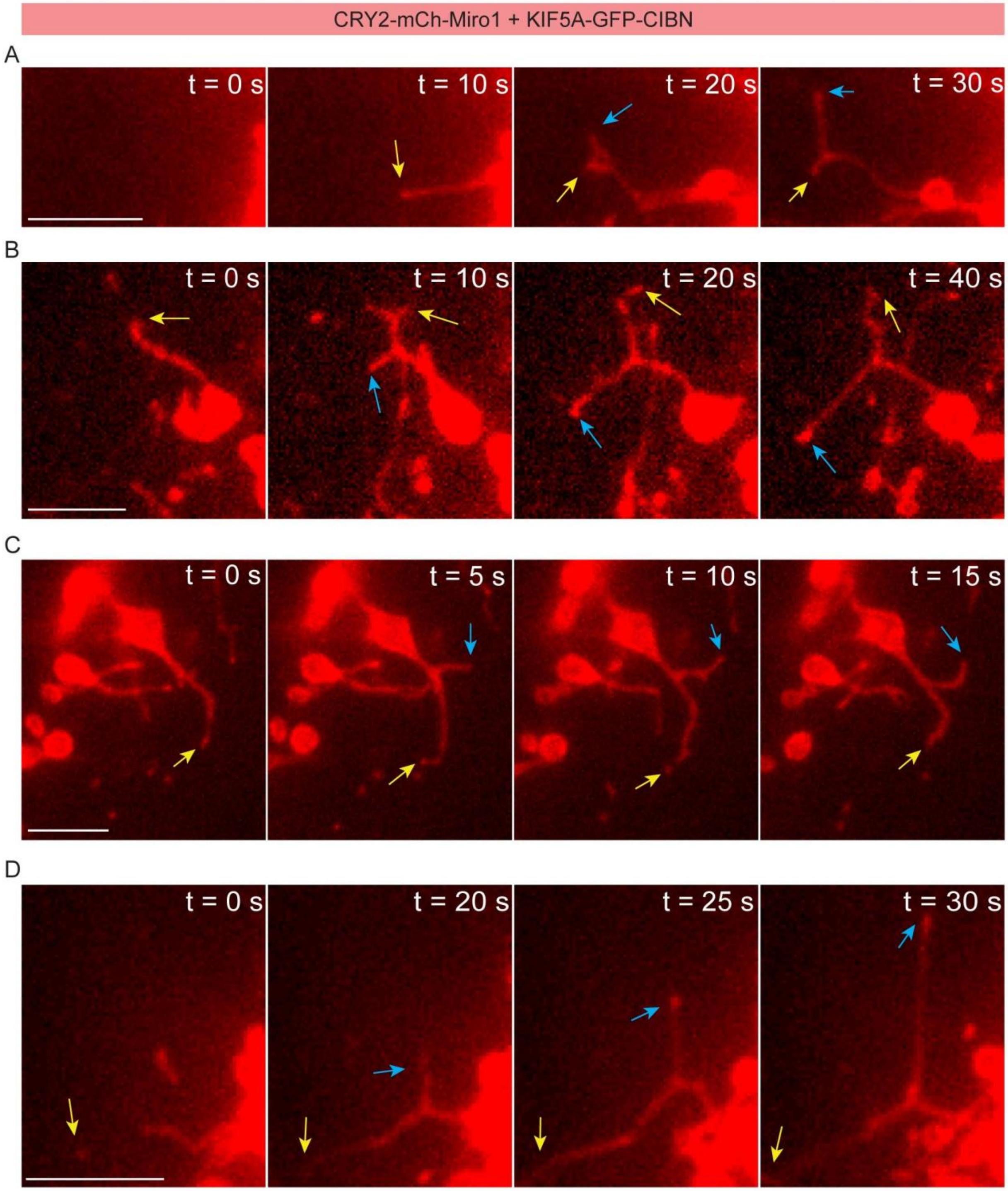


Figure S5. (A-D) Additional examples of mitochondrial outer membranes that show bifurcation of the stretching tubule after blue light activation. Scale bars, 5 μm.

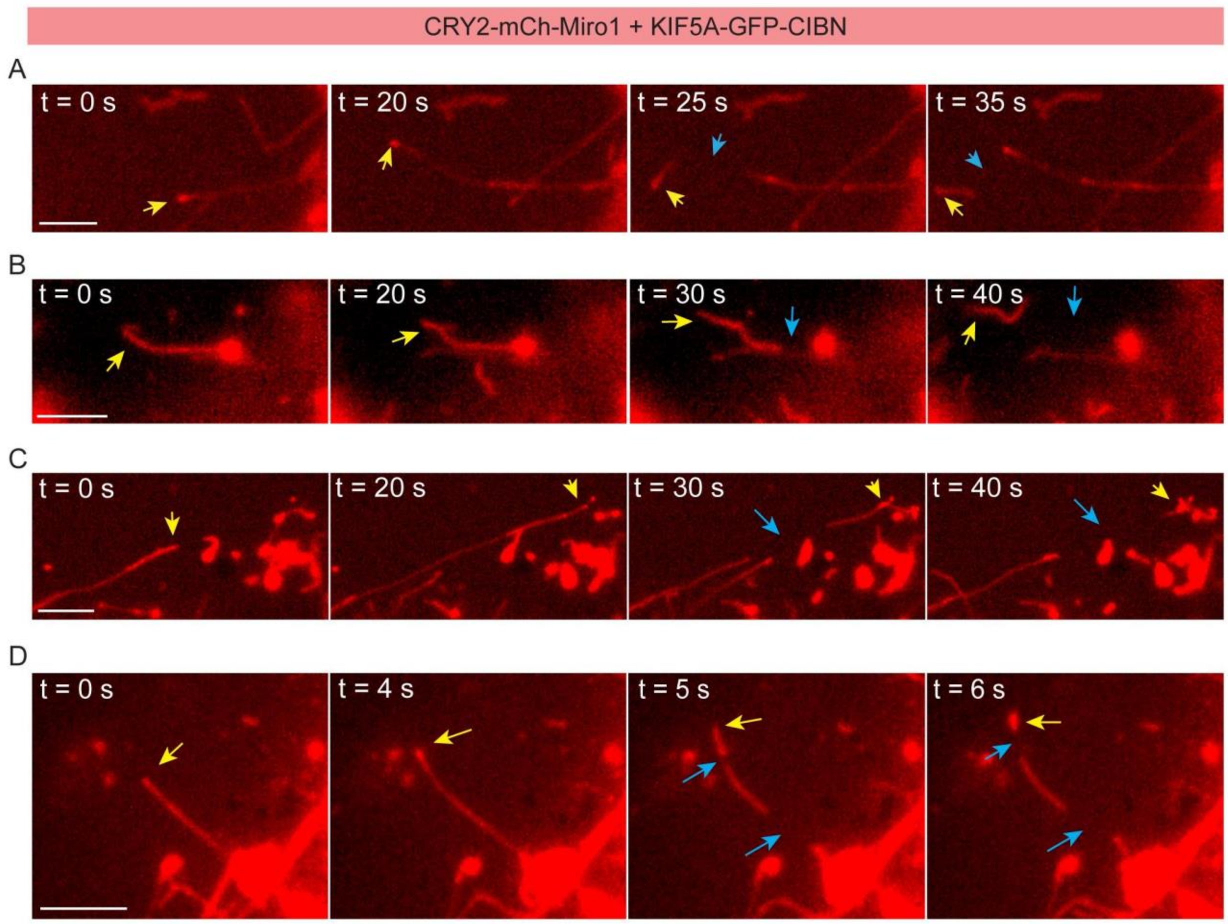


Figure S6. (A-D) Additional examples of mitochondrial outer membranes that show disconnection of the extending tubule after blue light activation. Scale bars, 5 μm.

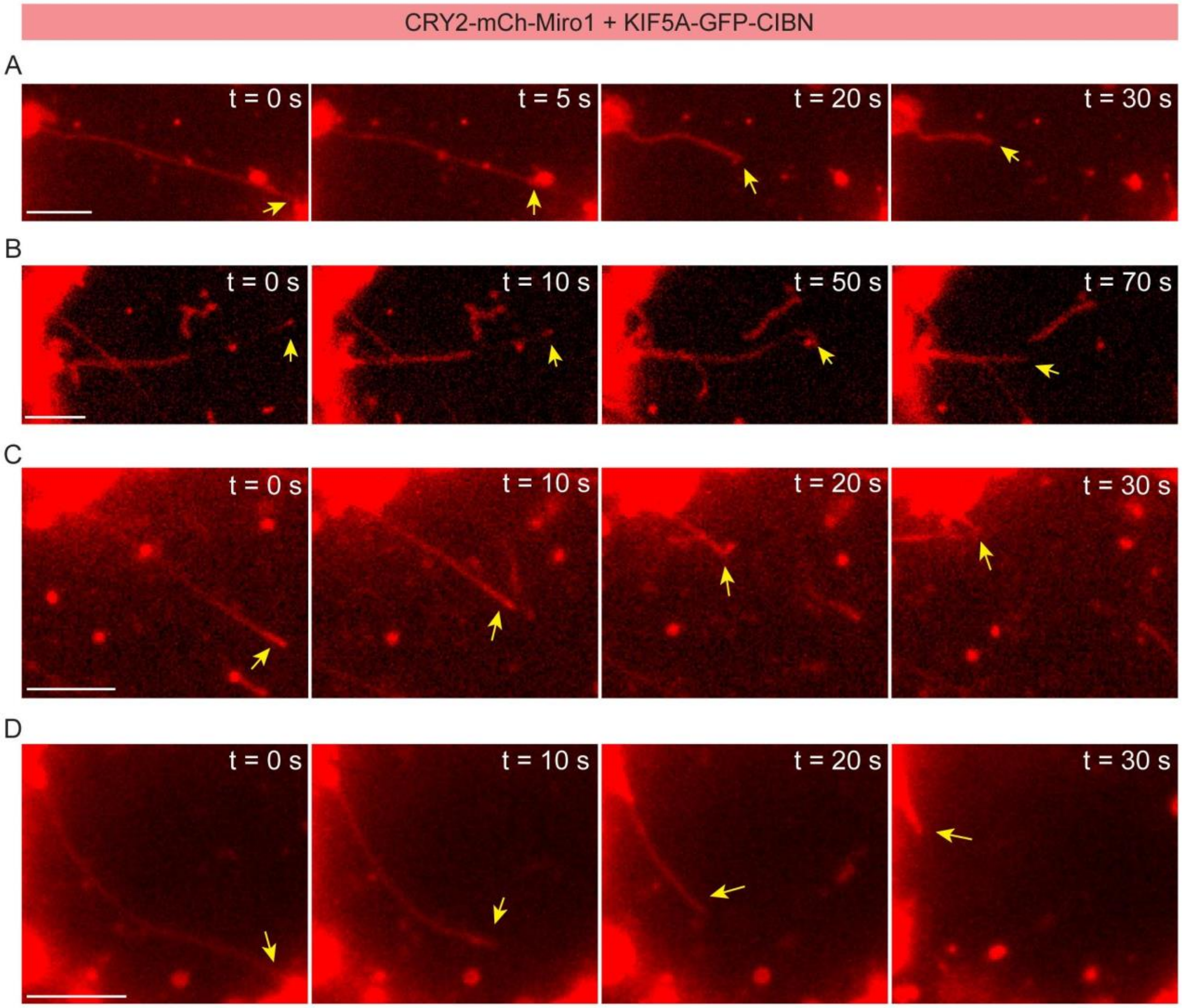


Figure S7. (A-D) Additional examples of mitochondrial outer membranes that show the retrieving of the extending tubule after blue light activation. Scale bars, 5 μm.

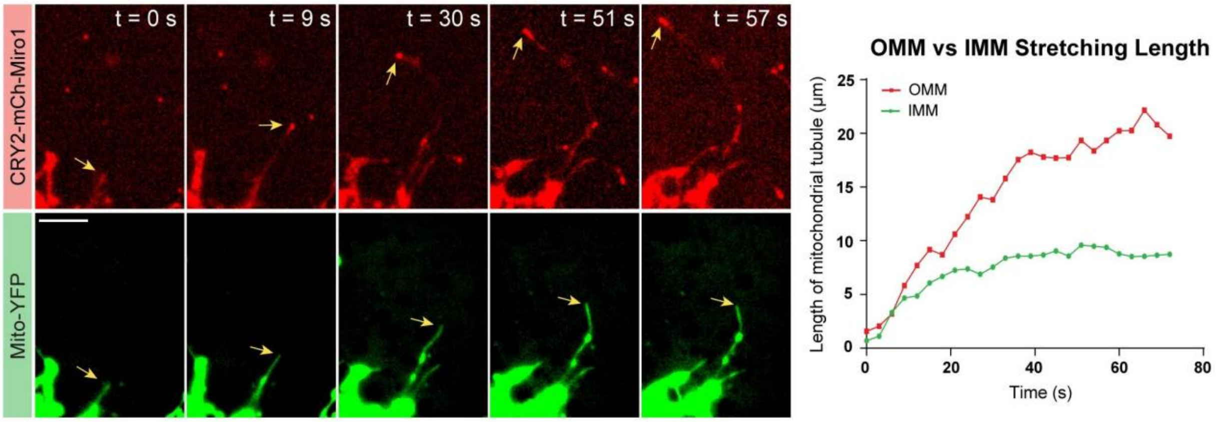


Figure S8. The separation between OMM and IMM can be significant. In the COS-7 cell transfected with CRY2-mCh-Miro1, KIF5A-CIBN and Mito-YFP, IMM tubule exhibits a significant lagging extension compared to the OMM tubule, as also revealed by the graph on the right showing lengths of extension for both tubules with respect to time. Scale bar, 5 μm.

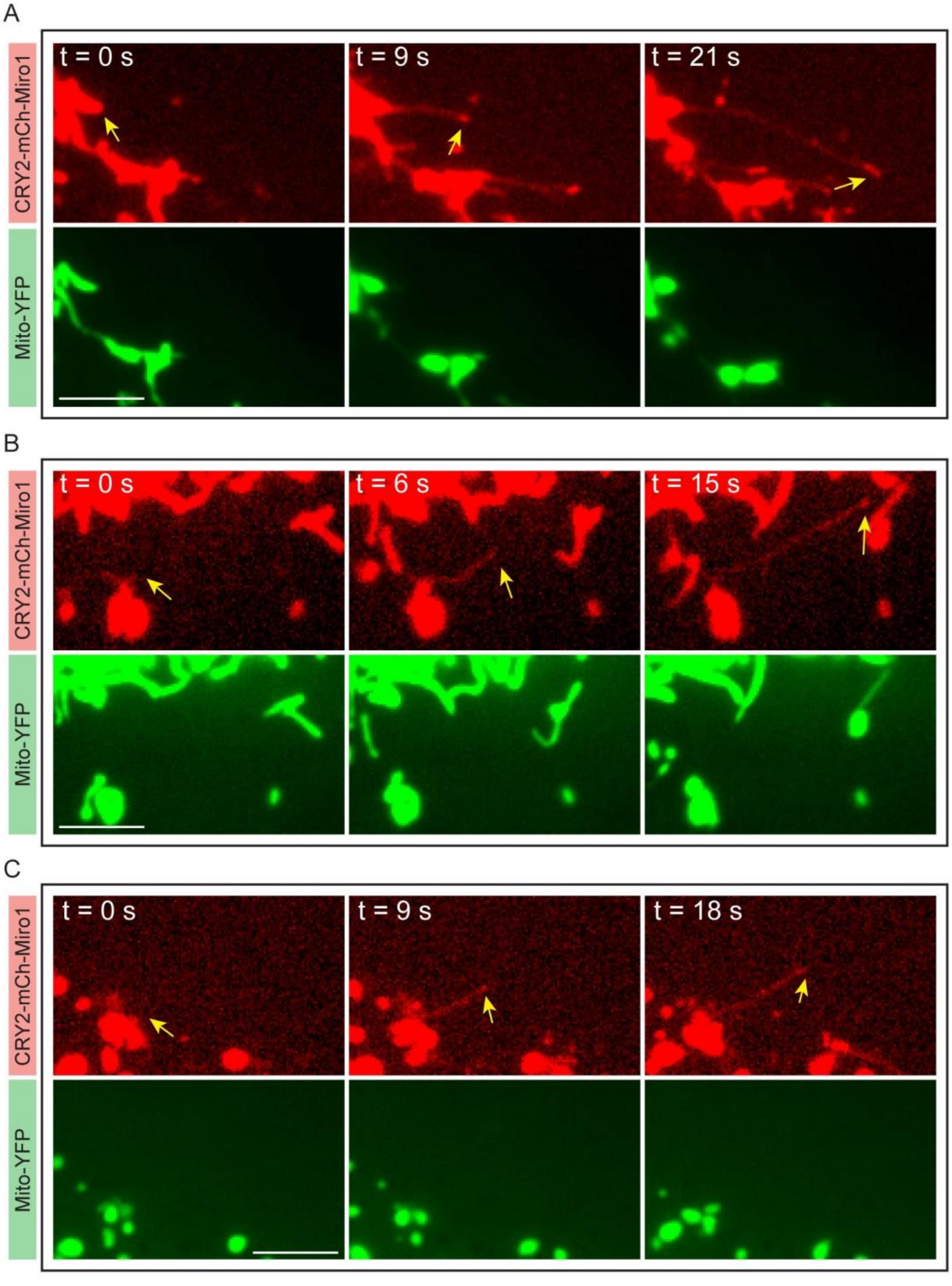


Figure S9. (A-C) Examples of mitochondrial inner membranes that do not show deformation after blue light activation although the outer membrane is deformed. Scale bars, 5 μm.

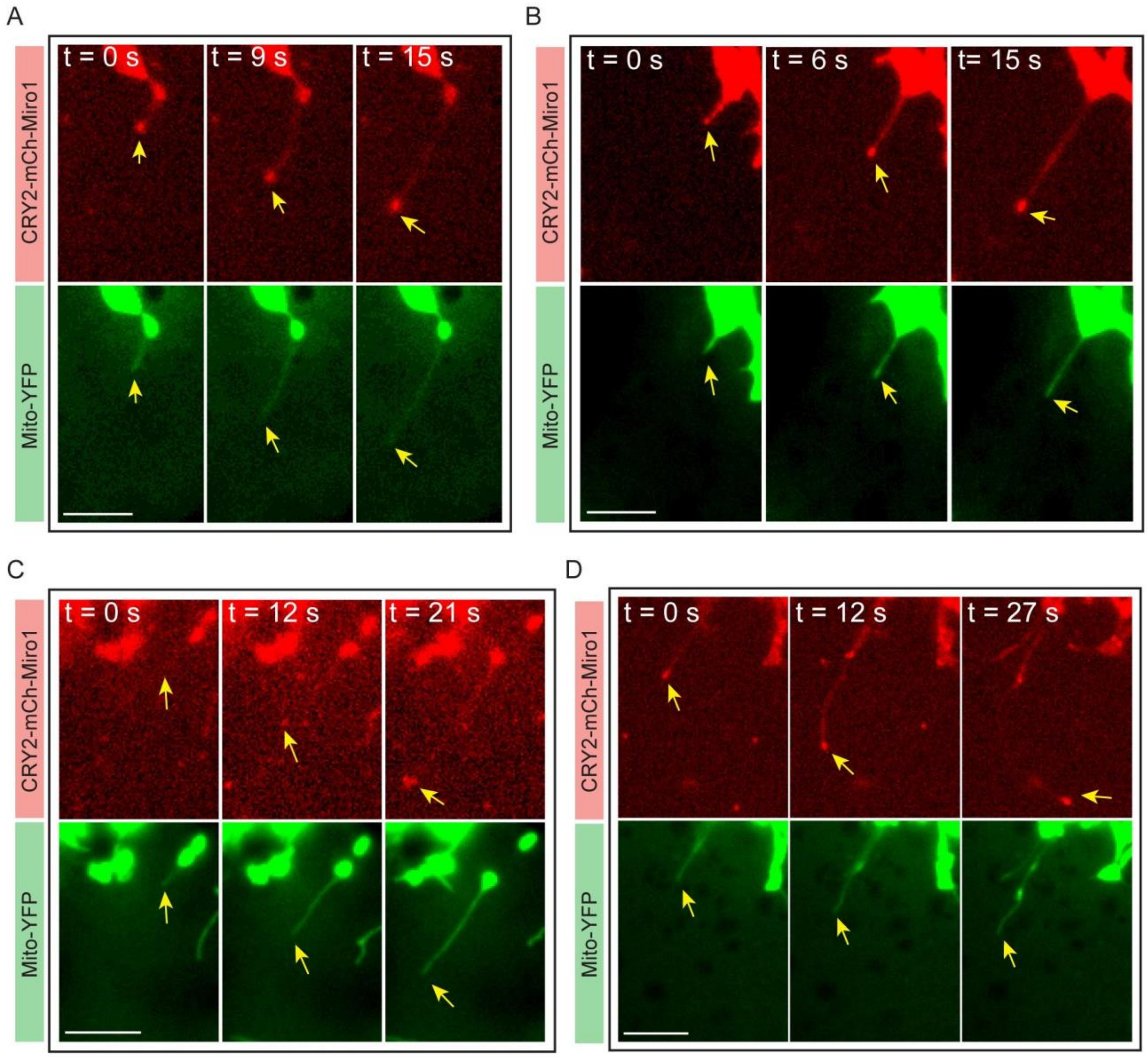


Figure S10. (A-D) Additional examples of deformation of mitochondrial inner membranes that show a tubule stretching out after blue light activation. Scale bars, 5 μm.

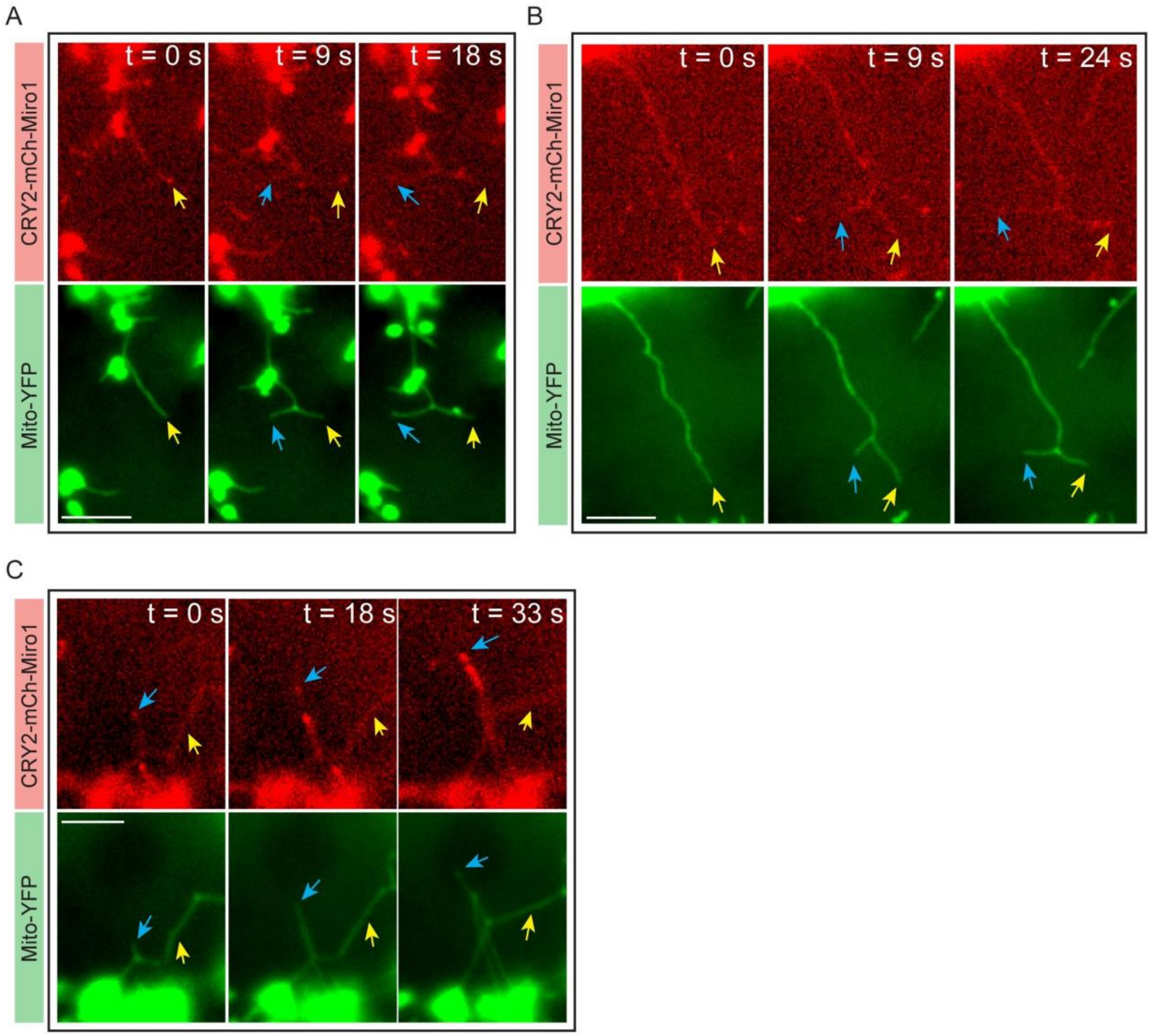


Figure S11. (A-C) Additional examples of mitochondrial inner membranes that show bifurcation of the stretching tubule after blue light activation Scale bars, 5 µm.

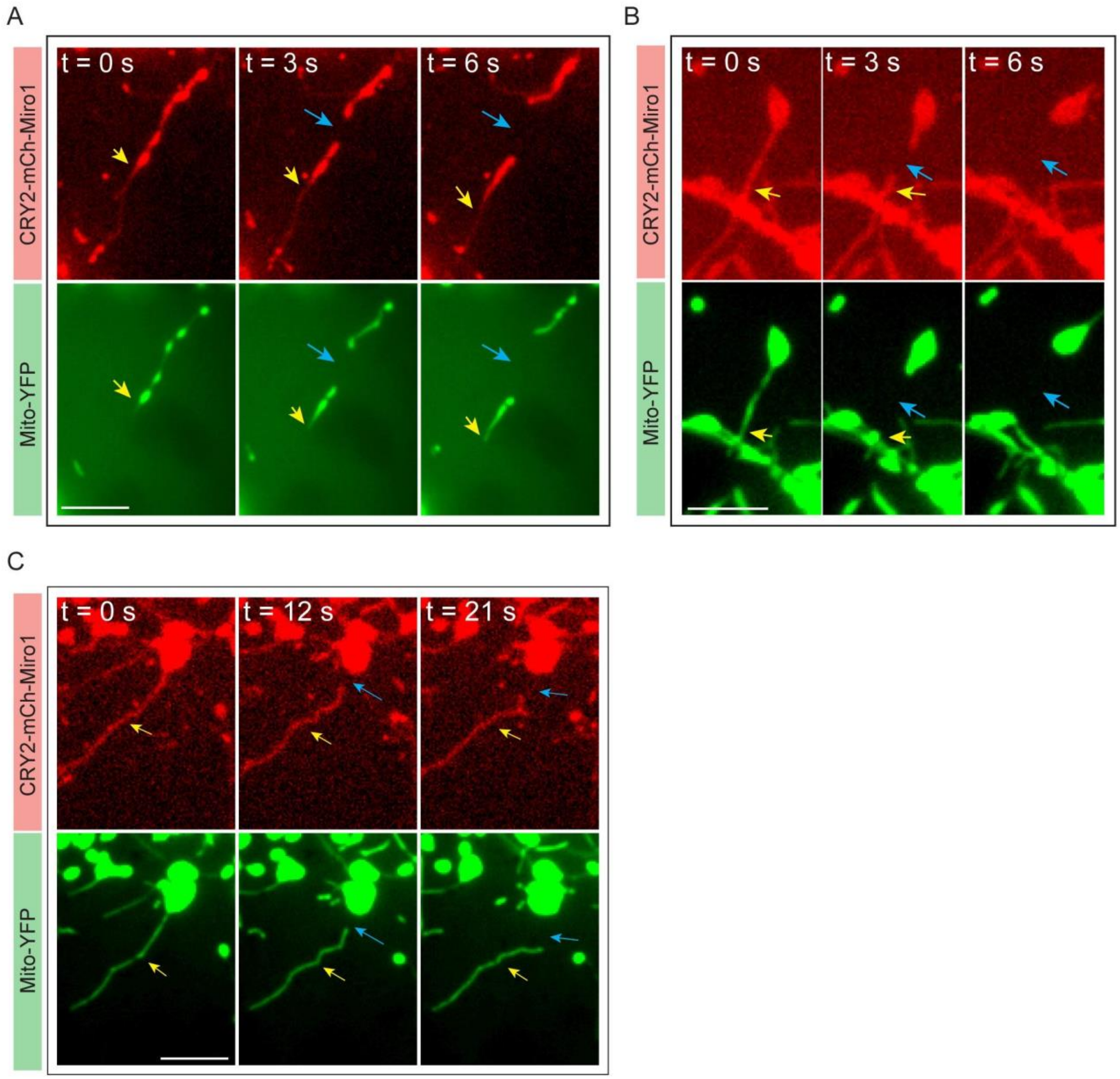


Figure S12. (A-C) Additional examples of mitochondrial inner membranes that show disconnection of the extending tubule after blue light activation. Scale bars, 5 μm.

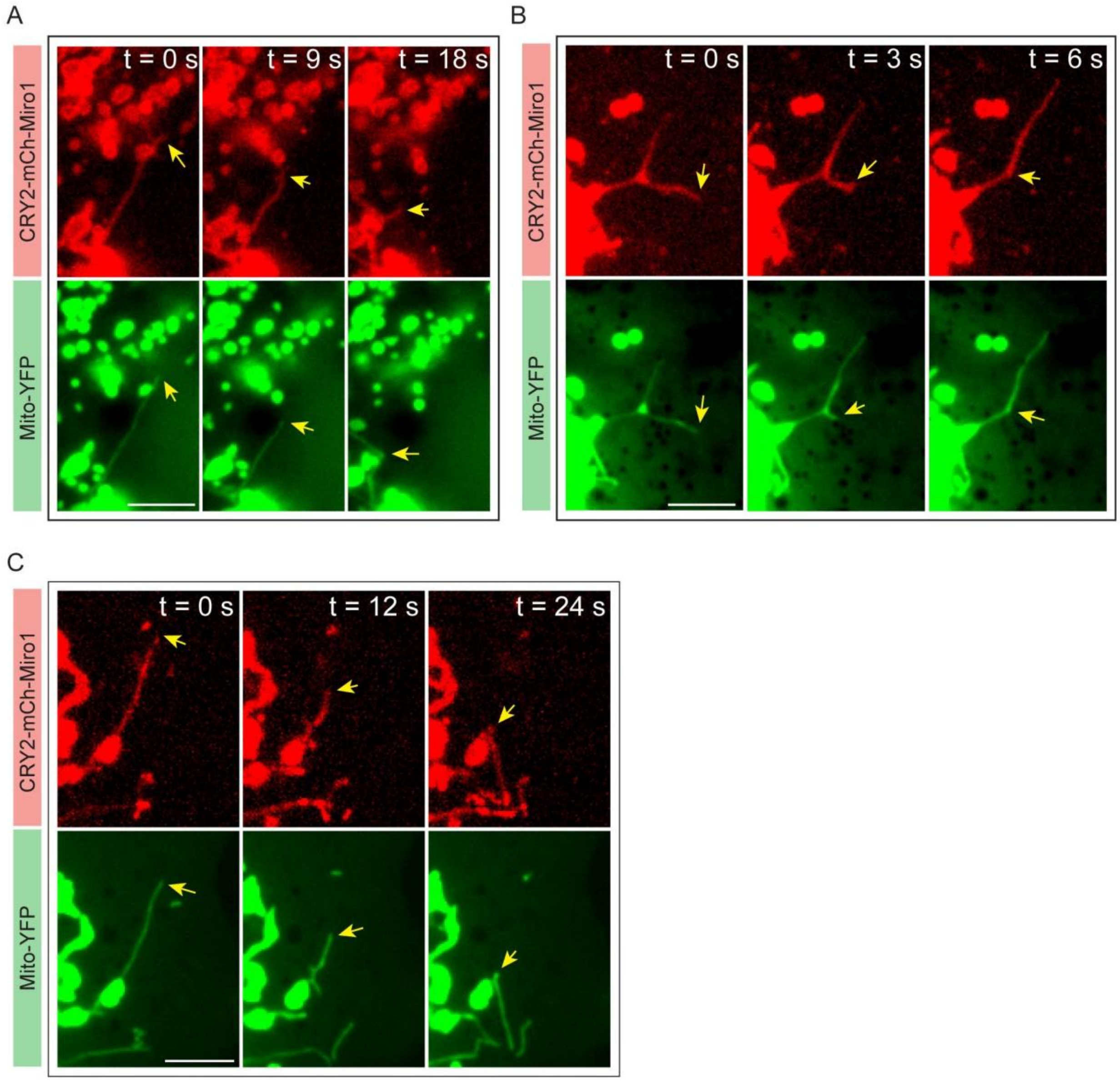


Figure S13. (A-C) Additional examples of mitochondrial inner membranes that show retrieving of the extending tubule after blue light activation. Scale bars, 5 μm.

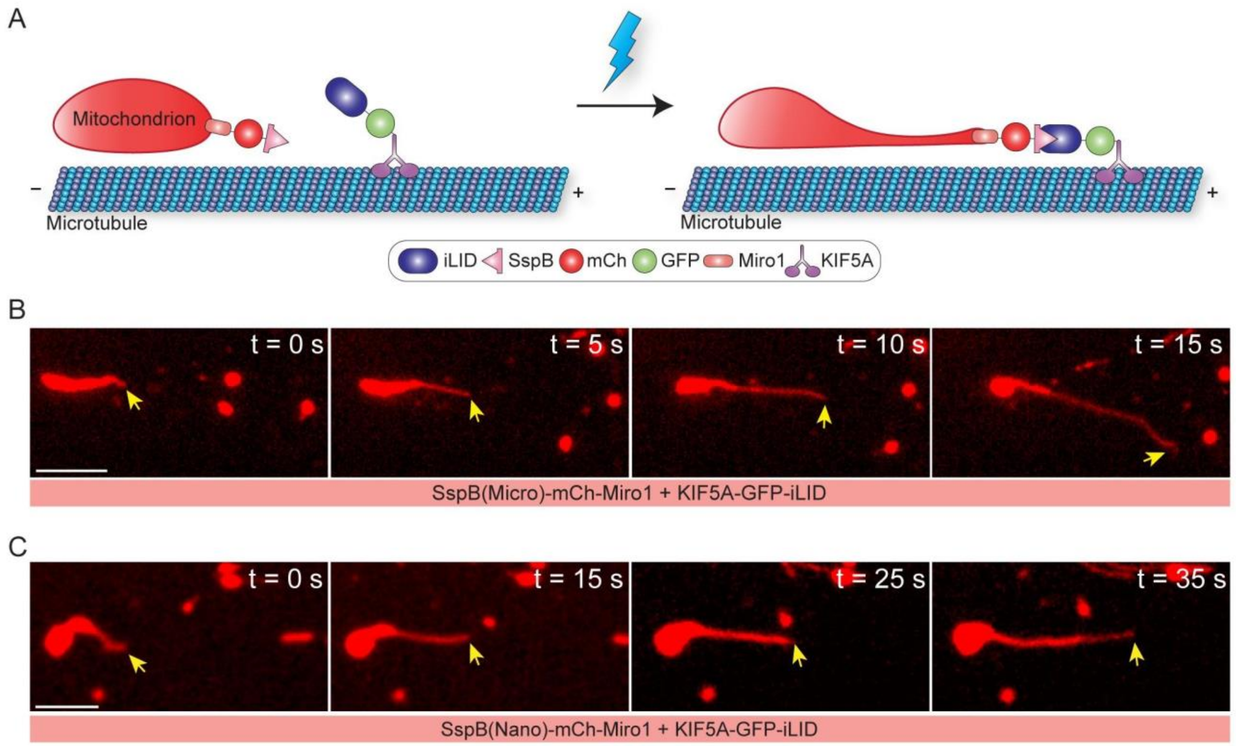


Figure S14. The iLID-SspB systems can also induce membrane deformation in mitochondria. (A) Scheme for another iLID-SspB system where SspB(Micro) is fused to Miro and iLID is fused to kinesin. (B) In the COS-7 cell transfected with SspB(micro)-mCh-Miro1 and KIF5A-GFP-iLid, deformation of mitochondria can also be induced, suggesting the exchange of positions of the pair in the two constructs does not affect the effectiveness of light-induced reshaping of mitochondria. (C) The integration of SspB(Nano), another variant of iLID binding partner, can also induce mitochondrial outer membrane deformation, in the COS-7 cell expressing SspB(Nano)-mCh-Miro1 and KIF5A-GFP-iLID. Scale bars, 5 μm.

**Movie S1.** Mitochondria showing deformation and elongation after blue light stimulation. Related to Figure 1B.

**Movie S2.** OMM in mitochondria shows light-induced stretching in the red channel. Simultaneously, the deformation and elongation of IMM tubules followed the same stretching paths of OMM tubules, as shown in the green channel. Related to Figure 3B.

### Tables

#### Table S1. Plasmids Constructions.

| Plasmids | Method | Template | Insertion sites | Forward Primer | Reverse Primer |
|---|---|---|---|---|---|
| Tau-GFP | Ligation | Tau-YFP | AgeI and NotI | - | - |
| iLID-mCh-Miro1 | In-Fusion | CIBN-mCh-Miro1 | NheI and AgeI | CGTCAGATCCGCTAGCGccaccGGGGAGTTTCTGGCAACCACac | CATGGTGGCGACCGGTccAAAGTAATTTTCGTCGTTCGC |
| KIF5A-GFP-SspB(Micro) | In-Fusion | KIF5A-GFP-CIBN | BSP1407I and AflII | ATGGACGAGCTGTACAagggcattgatctgagcggcctgac | CAATTTACGCcttaagTTAACCAATATTCAGCTCGTCATAG |
| SspB(Micro)-mCh-Miro1 | In-Fusion | CIBN-mCh-Miro1 | NheI and AgeI | CGTCAGATCCGCTAGCGccaccATGgAATTCAGCTCCCCGAAACGCc | ATGGTGGCGACCGGTccactaccACCAATATTCAGCTCGTCATAGATTTC |
| SspB(Nano)-mCh-Miro1 | In-Fusion | CIBN-mCh-Miro1 | NheI and AgeI | CGTCAGATCCGCTAGCGccaccATGgAATTCAGCTCCCCGAAACGCc | ATGGTGGCGACCGGTccactaccACCAATATTCAGCTCGTCATAGATTTC |
| KIF5A-GFP-iLID | In-Fusion | KIF5A-GFP-CIBN | BSP1407I and AflII | ATGGACGAGCTGTACAAGggtagtggtagtGGGGAGTTTC | CAATTTACGCcttaagttaAAAGTAATTTTCGTCGTTCGCTGC |
| KIF1A-GFP-SspB(Micro) | In-Fusion | KIF5A-GFP-SspB(Micro) | NheI and SalI | CGTCAGATCCgctagcgccaccatggctggggcctctgtgaag | GCTCAcCATAGTCGAcccggatcCcagcagatctcgcagcctgg |
| GFP-BICDN-SspB(Micro) | In-Fusion | GFP-CIBN | EcoRI and BamHI | ggtgggccgagaaTTCggcattgatctgagcggcctg | TAGATCCGGTGGATCCTTAACCAATATTCAGCTCGTC |
